# Genetic requirements for uropathogenic *E. coli* proliferation in the bladder cell infection cycle

**DOI:** 10.1101/2023.12.05.570251

**Authors:** Daniel G. Mediati, Tamika A. Blair, Ariana Costas, Leigh G. Monahan, Bill Söderström, Ian G. Charles, Iain G. Duggin

**Author notes:** Address correspondence to Iain Duggin.

## Abstract

Uropathogenic *Escherichia coli* (UPEC) requires an adaptable physiology to survive the wide range of environments experienced in the host, including gut and urinary tract surfaces. To identify UPEC genes required during intracellular infection, we developed a transposon-directed insertion-site sequencing (TraDIS) approach for cellular infection models and searched for genes in a library of ∼20,000 *E. coli* UTI89 transposon-insertion mutants that are specifically required for growth in M9-glycerol minimal medium, and at the distinct stages of infection of cultured bladder epithelial cells. Some of the functional requirements apparent for growth in M9-glycerol overlapped with those for the intracellular stage of infection, notably nutrient utilization, polysaccharide and macromolecule precursor biosynthesis, and cell envelope stress tolerance. Two genes implicated in both conditions were confirmed through independent gene deletion studies: *neuC* (sialic acid capsule biosynthesis) and *hisF* (histidine biosynthesis). Distinct sets of UPEC genes were also implicated in bacterial dispersal, where UPEC erupt from bladder cells in highly filamentous or motile forms upon exposure to human urine, and during recovery from infection in rich (LB) medium. Genes linked to septal peptidoglycan processes, *ytfB* and *dedD*, appeared to play roles in dispersal and may help stabilize cell division or the envelope during envelope stress created during infection. Our findings support a view that the host intracellular environment and infection cycle are multi-nutrient limited and create stress that demand an array of biosynthetic, cell envelope integrity and biofilm-related functions of UPEC.

**IMPORTANCE:** Urinary tract infections (UTIs) are one of the most frequent infections worldwide. Uropathogenic *Escherichia coli* (UPEC), which accounts for ∼80 % of UTIs, must rapidly adapt to highly variable host environments, such as the gut, bladder sub-surface and urine. In this study, we searched for UPEC genes required for bacterial growth and survival throughout the cellular infection cycle. Genes required for *de novo* synthesis of biomolecules and cell envelope integrity appeared to be important, and other genes were also implicated in bacterial dispersal and recovery from infection of cultured bladder cells. With further studies of individual gene function, their potential as therapeutic targets may be realized. This study expands knowledge of the UTI infection cycle and establishes an approach to genome-wide functional analyses of stage-resolved microbial infections.

## INTRODUCTION

Microbes are frequently challenged by highly variable environments and limited supplies of essential nutrients. An important example of the need to flexibly adapt to such dynamic conditions is provided by uropathogenic *Escherichia coli* (UPEC), which originate from the gut but have adapted the ability to disseminate and colonize the urinary tract. In an acute urinary tract infection (UTI), UPEC adhere to the luminal surface of the bladder epithelium and become internalized [1]. In the host cell cytoplasm UPEC proliferate and form biofilm-like intracellular bacterial communities (IBCs) [2]. When an IBC reaches an advanced state (up to 10^5^ bacteria per host cell), the IBC switches to a dispersal mode, accompanied by host cell rupture and the release of subpopulations of highly filamentous or motile UPEC [3, 4]. The diverse range of environments UPEC experience in the gut, urinary tract epithelium and urine—often involving rapid transitions—underscores the importance of a flexible metabolic capacity [5].

Genome-wide approaches to identify genes encoding the transporters, regulators, and metabolic enzymes required for growth under various culture conditions provide a way to understand the metabolic capacity and flexibility needed for infection and may lead to the identification of potential targets for antimicrobial therapies. Early studies assessed *E. coli* gene essentiality under rich and minimal nutrient media by screening collections (libraries) of transposon-insertion or knock-out mutant strains [6, 7]. More recently, techniques for mapping transposon-insertion libraries using high-throughput DNA sequencing, collectively known as transposon insertion-site sequencing (TIS), have expanded capacity for identifying essential genes [8]. The essential genes are identified by a lack of transposon insertions within those genes in a large transposon-insertion library pooled in one culture [9, 10]. The approach has also been developed to identify conditionally essential genes, such as after exposure to serum [11], bacteriophage [12], and antimicrobials [13–15], or through various physical selection methods that define cellular phenotypes or developmental pathways [8, 16].

Similar approaches to identify genes required for infection have been limited by the moderate scales possible with many experimental infections, combined with the large number of bacterial strains that would need to be simultaneously screened with TIS [8, 17]. Nevertheless, several studies have used TIS to map pathogen genes important for infection in suitable animal models by comparing the overall input and output pools of mutants [18–22]. TraDIS was first applied to UPEC to identify the gene profile required for in vitro growth of EC958 (an *E. coli* ST131 clonal lineage) in human serum [11]. Recently, UPEC gene requirements in acute UTI were assessed using pig and diuretic-mouse models, to help overcome the limited scale of the standard mouse model of cystitis [23, 24]. The results suggested that many bacterial genes, especially those involved in certain nutrient uptake and LPS biosynthesis pathways, are important in UTI.

Here we sought to identify genes specifically required during key stages of the intracellular infection cycle by using a stage-resolved cell culture model of infection [3, 25, 26]. We first modified the TraDIS protocol to streamline sequencing, with less input DNA required than the original protocol [10], and applied this to a moderate-scale transposon-insertion library suitable for infection models, consisting of ∼20,000 unique mutants in the model cystitis strain, *E. coli* UTI89. We modified and scaled-up the bladder epithelial cell infection model where feasible to make it more suitable for genetic screens. By mapping UPEC gene importance at the distinct stages of cellular infection, we found that many of the metabolic genes required in pure culture in minimal medium were also important for intracellular survival during infection. Novel sets of genes implicated in the bacterial dispersal and recovery stages of infection were also identified, which may represent targets for further analyses of the intracellular infection cycle and for potential therapeutic intervention in UTI.

## RESULTS

### Construction of a transposon mutant library in E. coli UTI89 and its characterization by a modified TraDIS protocol

Transposon mutagenesis of *E. coli* UTI89 was done by electroporation of the mini-Tn5 synthetic transposon, Ez-Tn5 <KAN-2>, assembled with modified Tn5 transposase (‘transposasomes’). Eleven electroporations yielded 97,500 kanamycin-resistant colony forming units (cfu). Colonies were resuspended, pooled, and then stored in frozen aliquots (OD_600_ = 10), which upon thawing contained 1.18 x 10^10^ cfu per mL. We developed a modified TraDIS protocol (**Supplementary Figure 1**) by initially fragmenting total DNA from the library with tagmentation, which requires much less input DNA than the original method and may be advantageous in infection models where bacterial yields are limited (**Supplementary Figure 1A**). The Tn-genome junctions were then amplified from both ends of the transposon separately (**Supplementary Figure 1B**), so that standard Illumina sequencing could be used to identify the insertion sites, whilst avoiding the need for custom primers or dark sequencing cycles (**Supplementary Figure 1C**). Sequence data analysis identified an average of 18,756 unique insertion sites in the chromosome and 464 in the plasmid, pUTI89. Tn-insertions were distributed throughout the genome, equating to an average Tn-insertion frequency of one per 270 bp in the chromosome and slightly greater in pUTI89 (**Figure 1A**, **Supplementary Table 1B**).

**Figure 1.**
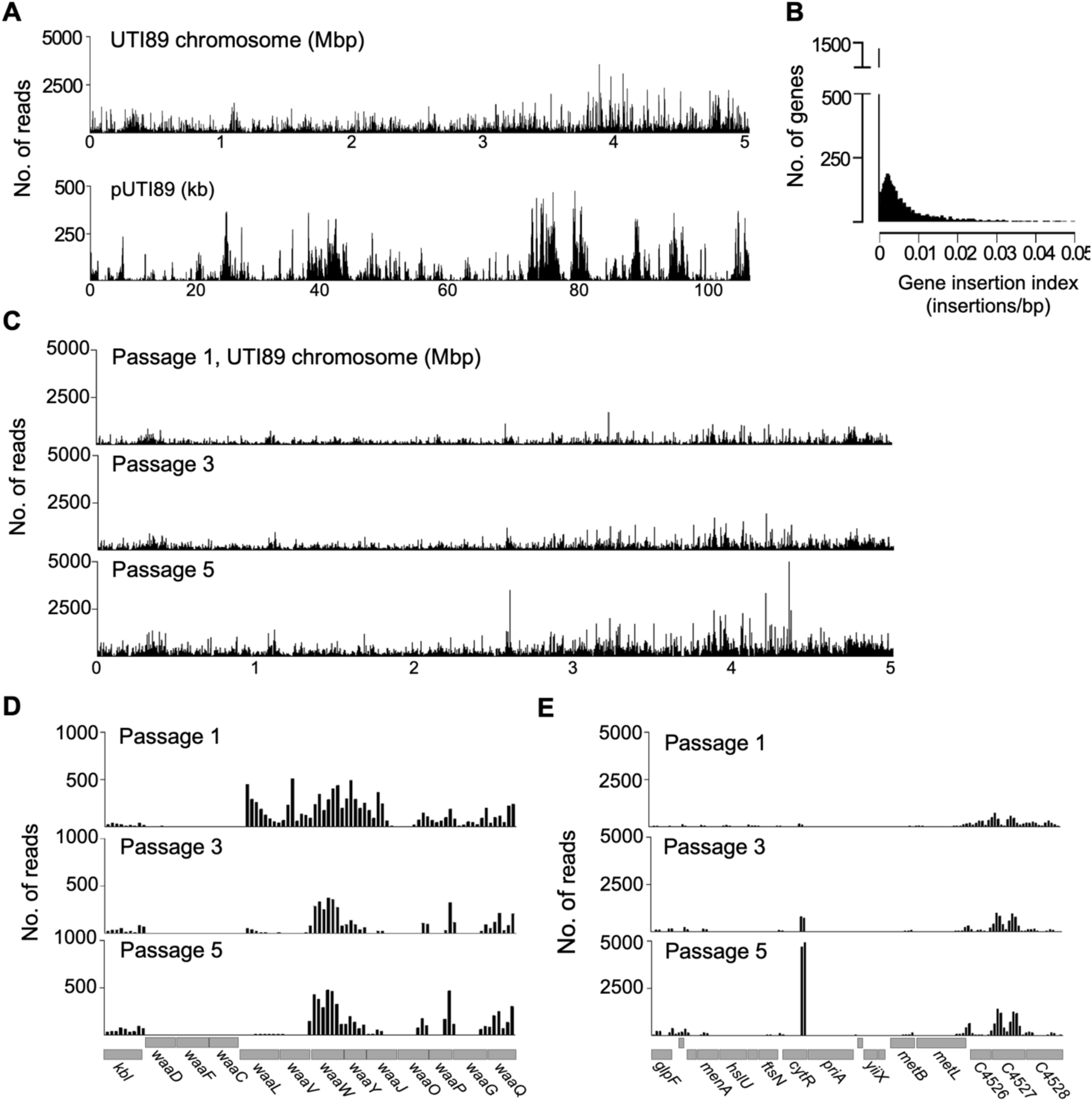
TraDIS mapping of an *E. coli* UTI89 transposon-insertion library. (A) The distribution of TraDIS mapped-reads in 150 bp windows along the 5.06 Mbp UTI89 chromosome and 110 kb plasmid pUTI89. (B) Frequency distribution of the gene insertion index—the number of Tn-insertions within each gene divided by that gene’s length (in bp)—in the UTI89 Tn-insertion library. (C) The number of mapped-reads in 150 bp windows along the UTI89 chromosome following daily passaging (1/400) in LB medium. Data were combined from both Tn-ends to generate these plots. (D) Examples of the read-count distribution (150 bp windows) from one culture replicate, encompassing 4,053,040 – 4,066,540 bp, showing a decrease in abundance of insertion mutants in *waaV* and *waaL*, and (E) 4,394,185 – 4,413,535 bp, showing an increase in abundance of insertion mutants in *cytR* over time.

A strong correlation (R^2^ = 0.878) was observed for the number of insertion sites identified within each annotated open reading frame (ORF) obtained from the two Tn-ends (**Supplementary Figure 2**). Approximately 22 % of genes (∼1,200) did not have a detected Tn-insertion (*i.e.*, they had an insertion index of zero, **Figure 1B**) and they are listed in **Supplementary Table 1E**. It has been predicted that 8-10 % of an *E. coli* genome contains genes that are individually essential for growth in LB medium [9, 27]. Many of the Tn-free genes in the UTI89 library are likely to be essential for viability, but in a library of this size a minor proportion of non-essential genes should also be Tn-free by chance alone. Consistent with this, the mean size of the Tn-free ORFs was 569 bp compared to the genome-wide mean of 901 bp. Genes containing at least one Tn-insertion gave rise to an approximately symmetrical frequency distribution of the pooled gene insertion index (insertions/gene/bp, **Figure 1B**), consistent with a near-random insertion frequency in these genes. The shoulder of this peak was not completely distinct from zero, further supporting the expectation that Tn-insertions are not present in all non-essential genes in this library [10, 28]. As such, not all genes will be tested in screens with this library, however it should avoid experimental bottlenecks—random loss of mutants caused by the insufficient feasible scale of the infection model.

To prepare for comparing the UTI89 Tn-library in different conditions including infection, we first did a preliminary analysis of the relative abundance of mutants in the library over time, by passaging the library with a 1/400 dilution in LB once per day over five days. Samples taken after day one, three and five were analysed by TraDIS (Figure 1C). Genome-wide analysis identified 196 genes with a substantial reduction in read counts from days one to five (log_2_ fold-change (FC) ≤ -2 and *p*-value ≤ 0.05), suggesting that disruption of these genes reduces the growth rate in LB (**Supplementary Table 1F**, *e.g.*, *waaL*, Figure 1D). Conversely, 16 genes showed a substantial increase in the relative abundance of Tn-insertion reads by day-five (**Supplementary Table 1F**, *e.g.*, *cytR*, Figure 1E). These results reflect the dynamic behaviour of transposon mutant libraries and may serve as a reference for comparing changes seen in other conditions over time, to improve the specificity of identifying conditionally essential genes.

### Identification of genes important for UTI89 growth in M9-glycerol

To identify genes that become important for growth or survival during a transition to nutrient-limited conditions, approximating nutritional changes that bacteria may experience when moving between various host environments, we diluted the stock library in LB then diluted 1/10 into M9-glycerol minimal medium and LB (control), in duplicate, and then sampled the four cultures after three 24 h passages in M9-glycerol (1/10 dilutions) for TraDIS. Read counts and basic statistics for all datasets are in **Supplementary Table 1B**, and the complete output from analyses comparing individual genes between M9-glycerol and LB are in **Supplementary Table 1G**. A total of 488 genes showed a difference (*p*-value ≤ 0.05) between these media in either one or both replicates, but more stringent criteria (*i.e.*, *p*-value ≤ 0.05 and log_2_FC ≤ -1, in both replicates) resulted in a list of 60 candidate genes important for growth in M9-glycerol and not in LB (Figure 2A). Many of the most significantly underrepresented mutants after transition to M9-glycerol (log_2_FC < -2) are implicated in catabolic functions and cell envelope integrity under stress (**Table 1**).

**Figure 2.**
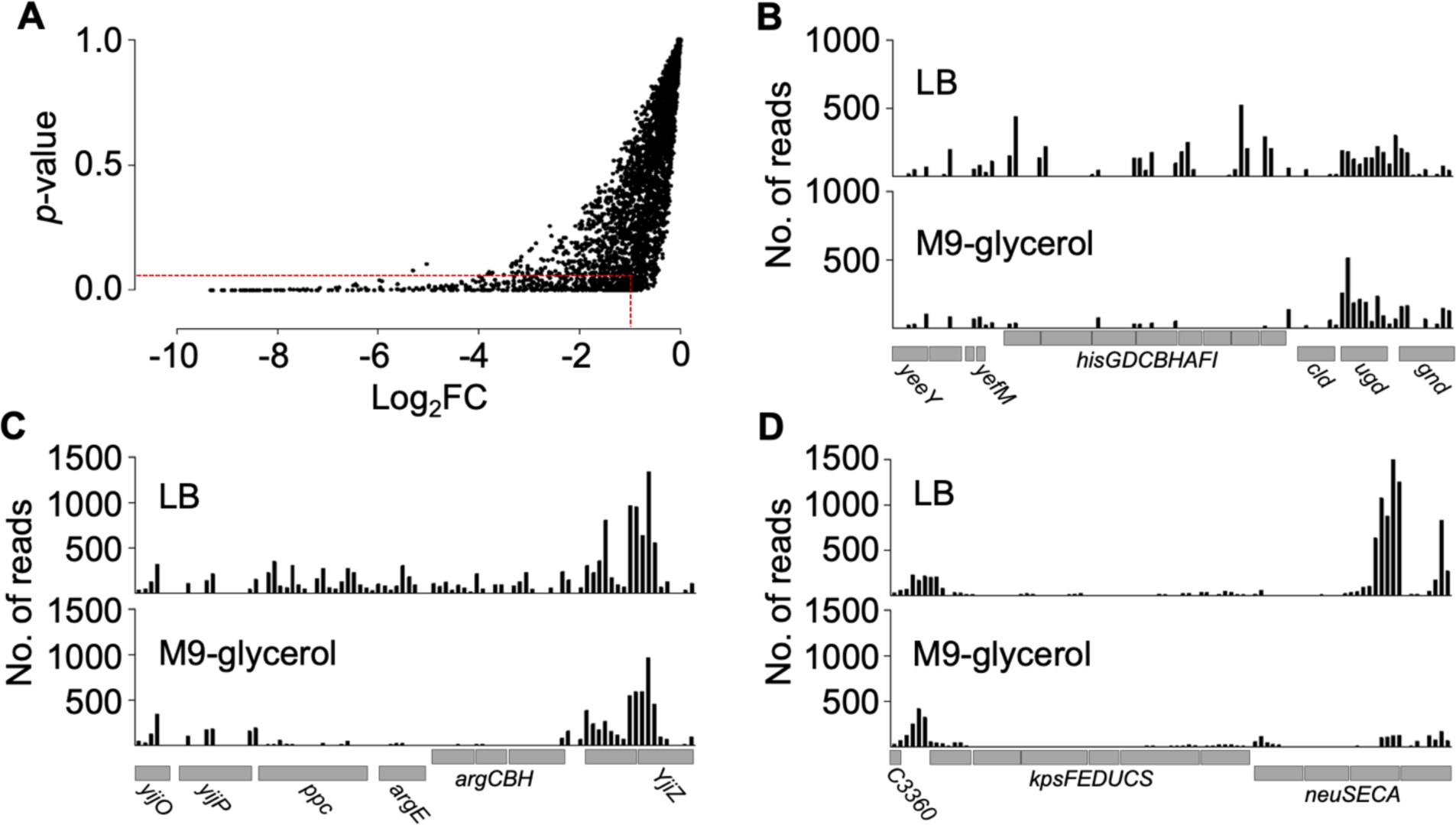
Identification of genes important for growth in M9-glycerol compared to LB. (A) Plot of each gene’s *p*-value against its corresponding log_2_FC value, resulting from the comparison of each gene’s Tn-insertion read count in M9-glycerol and LB media. The dashed lines indicate the stringent threshold criteria (*p*-value ≤ 0.05 and log_2_FC < -1). (B-D) Examples of frequency distributions of read counts from one biological replicate at selected loci (read count determined in 150 bp windows). The position of the annotated genes is indicated. (B) Region 2,225,678 – 2,239,628 bp. (C) Region 4,431,758 – 4,445,258 bp. (D) Region 3,288,114 – 3,301,914 bp.

**Table 1.**
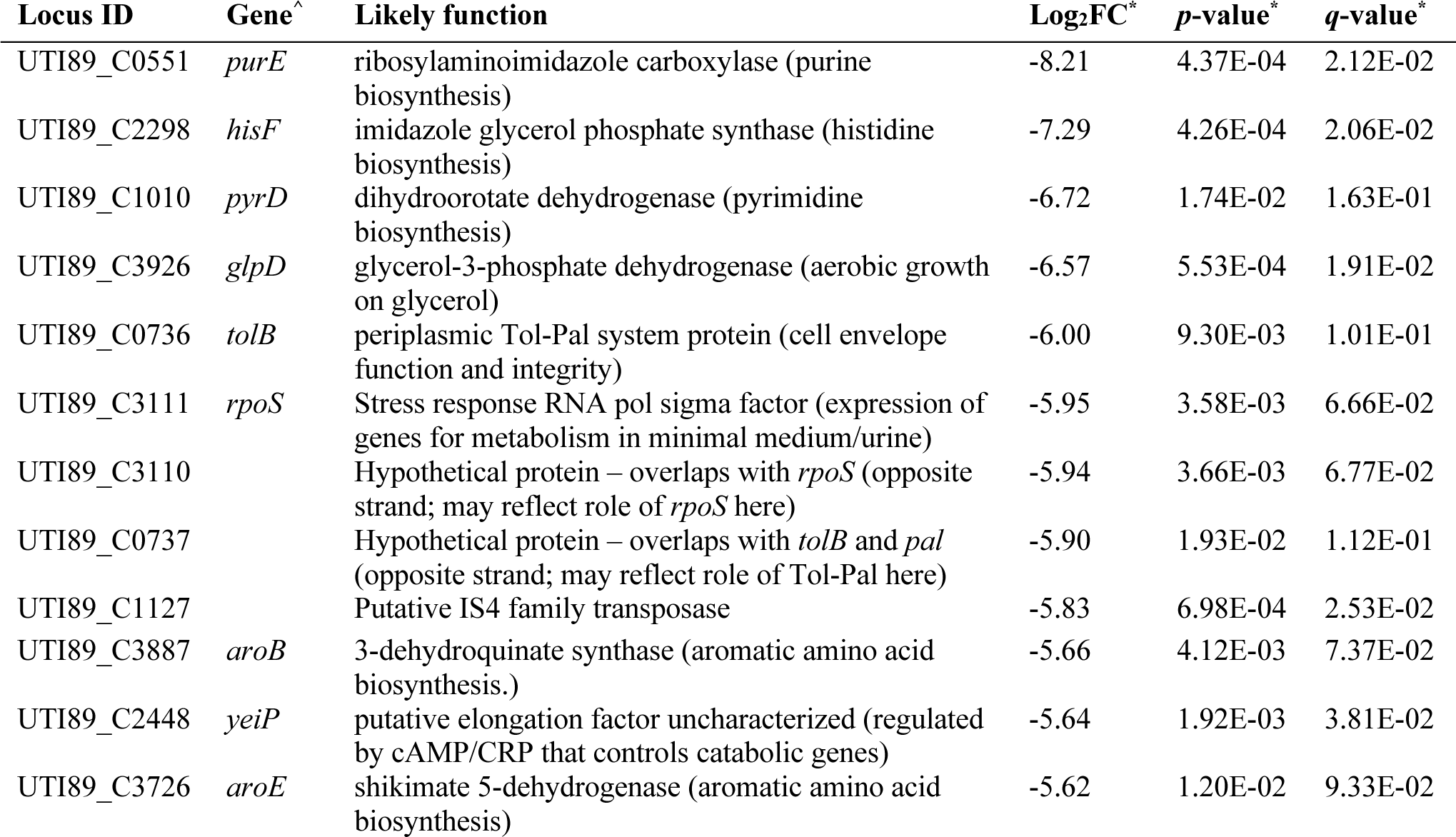
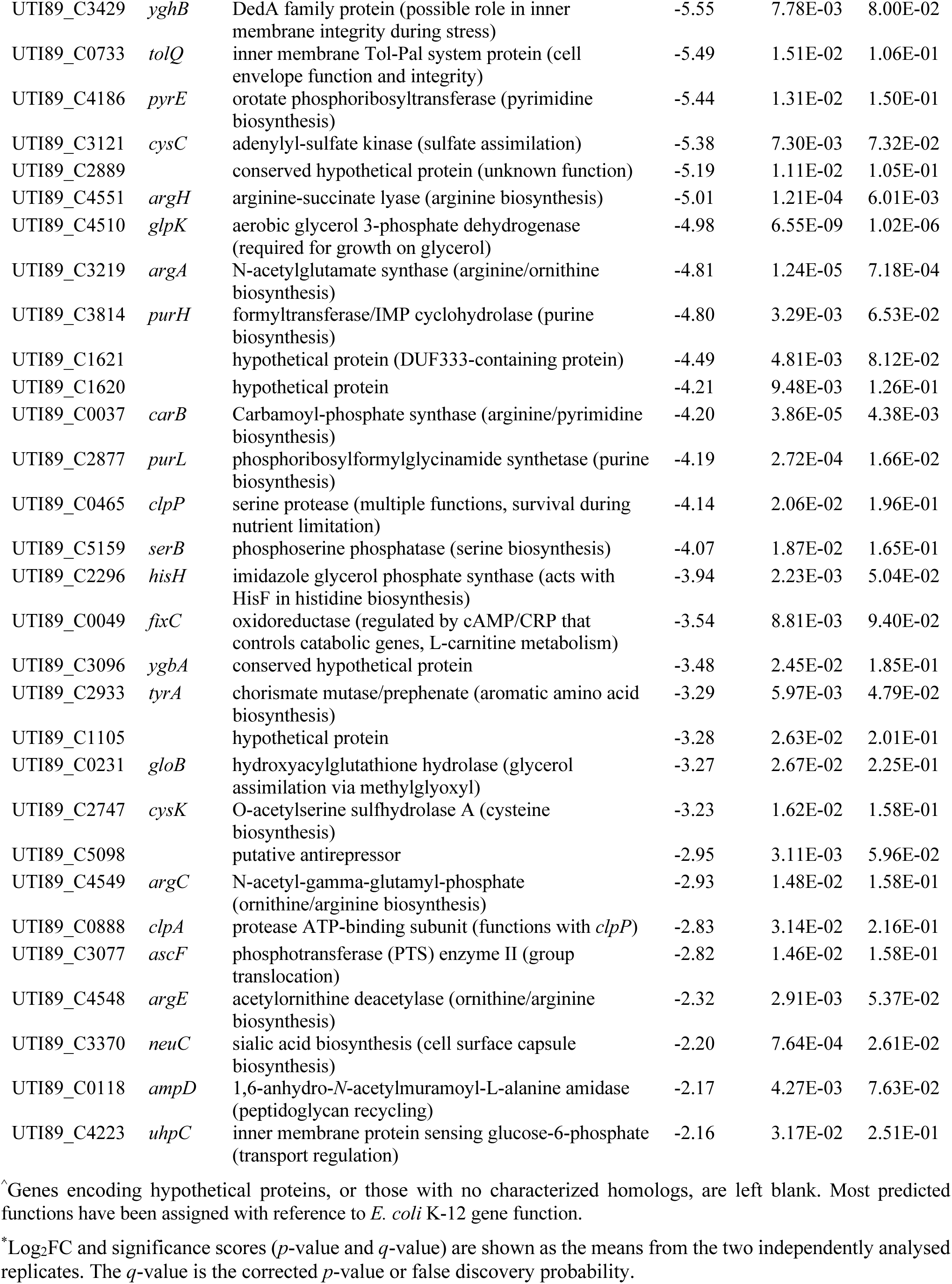
*E. coli* UTI89 candidate genes important for growth in M9-glycerol, compared to LB.

TraDIS data for three selected genomic regions containing significant differences between M9-glycerol and LB are shown in Figure 2B-D. These included the histidine biosynthesis operon (Figure 2B) and several genes for arginine biosynthesis (Figure 2C). These genes are required during nutrient-limited conditions in *E. coli* K-12, and many of the other genes important in UTI89 (**Table 1**) also have known roles in catabolic functions in M9-glycerol cultures of the commensal model strain K-12 [7]. A different example is *neuC* (Figure 2D), encoding an epimerase for sialic acid biosynthesis, expected to be involved in cell surface polysialic acid production. Interestingly, *neuC* and the gene cluster it sits within for capsule production are not present in many other strains of *E. coli* that can grow well in M9-glycerol (*e.g.*, K-12), consistent with the concept that gene requirements are strain and context dependent [29].

### Individual testing of selected UTI89 genes’ importance for growth in M9-glycerol

We selected five UTI89 genes that showed TraDIS results of differing effect size and statistical strength: *hisF*, *neuC*, *yggB*, *pdhR* and *ykgC* (**Table 1**, **Supplementary Table 1G**). These five genes were deleted from UTI89 using λ-Red recombination and then growth was monitored in M9-glycerol and LB to obtain the mean log-phase growth rates and maximal optical densities (OD_600_) as a percentage of wild-type (WT) (Figure 3, with growth curves shown in **Supplementary Figure 3**). In all five mutant strains, growth in LB was indistinguishable from WT UTI89. However, after transition to M9-glycerol their responses differed. The deletion of *hisF* resulted in complete inhibition of growth, consistent with the very significant difference between M9-glycerol and LB seen in TraDIS (**Table 1**). Deletion of *neuC* or *pdhR* resulted in a strong growth defect in M9-glycerol (Figure 3A, **3B**, and **Supplementary Figure 3**). This was expected, since the *pdhR* gene is also essential for growth of *E. coli* K-12 in M9-glycerol [7], although the *pdhR* results had not met the high-stringency thresholds applied to the UTI89 TraDIS data (see **Supplementary Table 1G**). Deletion of *yggB* in UTI89 resulted in only a moderate growth defect in M9-glycerol and showed a moderate reduction in the TraDIS (mean Log_2_FC = -1.69, *p* = 0.0002362). Lastly, deletion of *ykgC* did not affect growth in M9-glycerol (Figure 3), consistent with the TraDIS screen that showed only a minor fold-change between M9-glycerol and LB that varied between replicates (log_2_FC of -0.64 and -0.11, with *p*-values of 0.00779 and 0.706, respectively).

**Figure 3.**
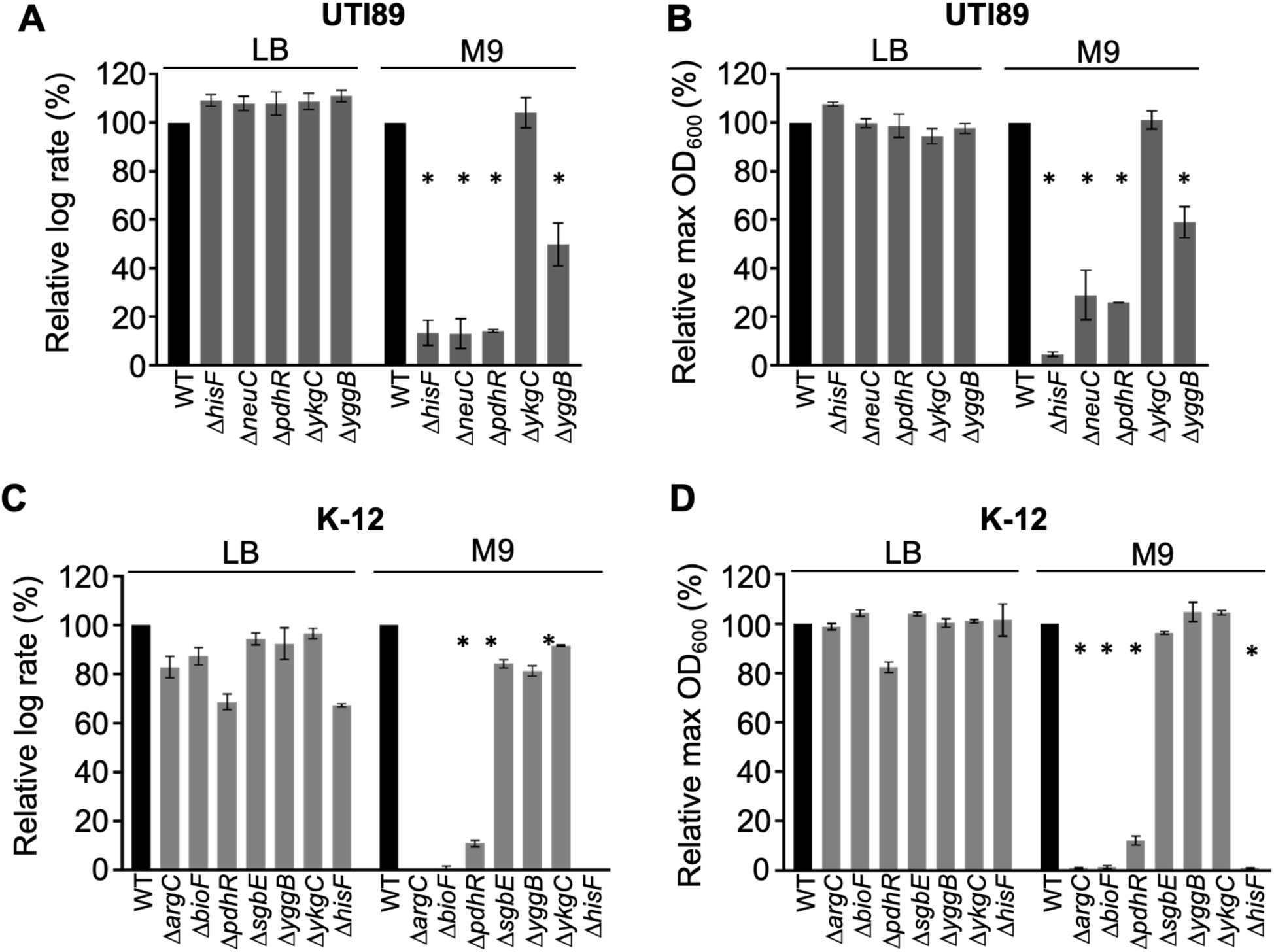
Growth of selected *E. coli* gene deletion mutants. (A) The relative log growth rates (µ) determined from growth curves (normalized as a percentage of WT UTI89) grown in LB or M9-glycerol. (B) The maximum OD_600_ (% of WT) over the 24 h growth period. Panels (C) and (D) represent the same approach applied to *E. coli* K-12 strain BW25113 and the indicated mutants. Error bars indicate the standard deviation (SD) from *n*=2 independent replicates. \**p*-value ≤ 0.05 (two-sided t-test compared to WT).

We next aimed to determine if some of the UTI89 results above were congruent with findings from a previous screen of *E. coli* K-12 gene knockouts in M9-glycerol [7]. We analyzed LB and M9-glycerol growth curves that we performed on seven K-12 strains from the Keio collection of knockout mutants [27]. Four of the mutant genes (*hisF*, *yggB*, *pdhR* and *ykgC*) are the direct homologs of genes analysed in UTI89 above, whereas the additional three (*argC*, *bioF*, and *sgbE*) were chosen to assess their correspondence with the range statistical strengths that their homologs exhibited in the UTI89 TraDIS analyses (**Table 1** and **Supplementary Table 1G**). Consistent with the UTI89 TraDIS results, the K-12 *argC*, *bioF* and *hisF* knockout strains grew well in LB but failed to show any significant growth in M9-glycerol (Figure 3C, **3D**, and **Supplementary Figure 4**). The K-12 *pdhR* knockout also grew very poorly in M9-glycerol as expected [7] although, beyond 24 h, eventually grew to high cell densities like the WT. The K-12 *yggB*, *ykgC* and *sgbE* deletions resulted in little or no growth defects in M9-glycerol, consistent with results obtained in UTI89. The growth results for these strains in UTI89 and K-12 therefore show a similar overall growth profile except for *neuC*, which is not found in K-12 yet has an important role for growth of UTI89 in M9-glycerol. These results support the TraDIS screen results, and the stringency of the thresholds we applied. However, the imperfect correspondence between TraDIS and individual tests in some cases highlights the value of independently testing and verifying of the role of genes based on the TraDIS screen results.

### Assessing UTI89 gene requirements during infection of cultured bladder epithelial cells

The UPEC intracellular infection cycle was first defined in infected mouse bladder explants using time-lapse microscopy [4]. Much remains unknown of the UPEC genetic requirements during each distinct stage of the acute infection cycle. One of the challenges is that the cellular infection stages resolvable by microscopy are quite asynchronous at the organ level so would not be resolved by transposon library screens on current *in vivo* infection models. Furthermore, the limited scale of standard *in vivo* infection models can also severely limit the simultaneous screening of large libraries of mutants (termed ‘bottlenecking’).

To begin addressing this challenge with TraDIS-based genetic screens, we developed a more scalable version of the established *in vitro* human bladder epithelial cell (BEC) infection model (see Methods), which enables sampling at the distinct stages of the intracellular infection cycle and recapitulates the stages observed by microscopy *in vivo* [4, 25, 30]. We first grew the UTI89 Tn-library statically in LB, then took a pre-infection (or ‘inoculate’) sample for TraDIS, and we infected three 100 mm culture dishes containing confluent BECs for each infection stage to be sampled (Figure 4A). Bacteria surviving intracellular infection were harvested at 24 h post-infection (IBC/intracellular infection stage) and 34-44 h post-infection (extracellular dispersal stage, urine exposure). Also, bacteria from a dispersal sample (34-44 h PI) were resuspended in LB and harvested after 18 h recovery, and a sample of this was subjected to a further 24 h passage in LB (42 h) (Figure 4A). Figure 4B-D show the relative bacterial yields over time and the appearance of the characteristic filamentous UPEC obtained from the dispersal stage of UTI with this model.

**Figure 4.**
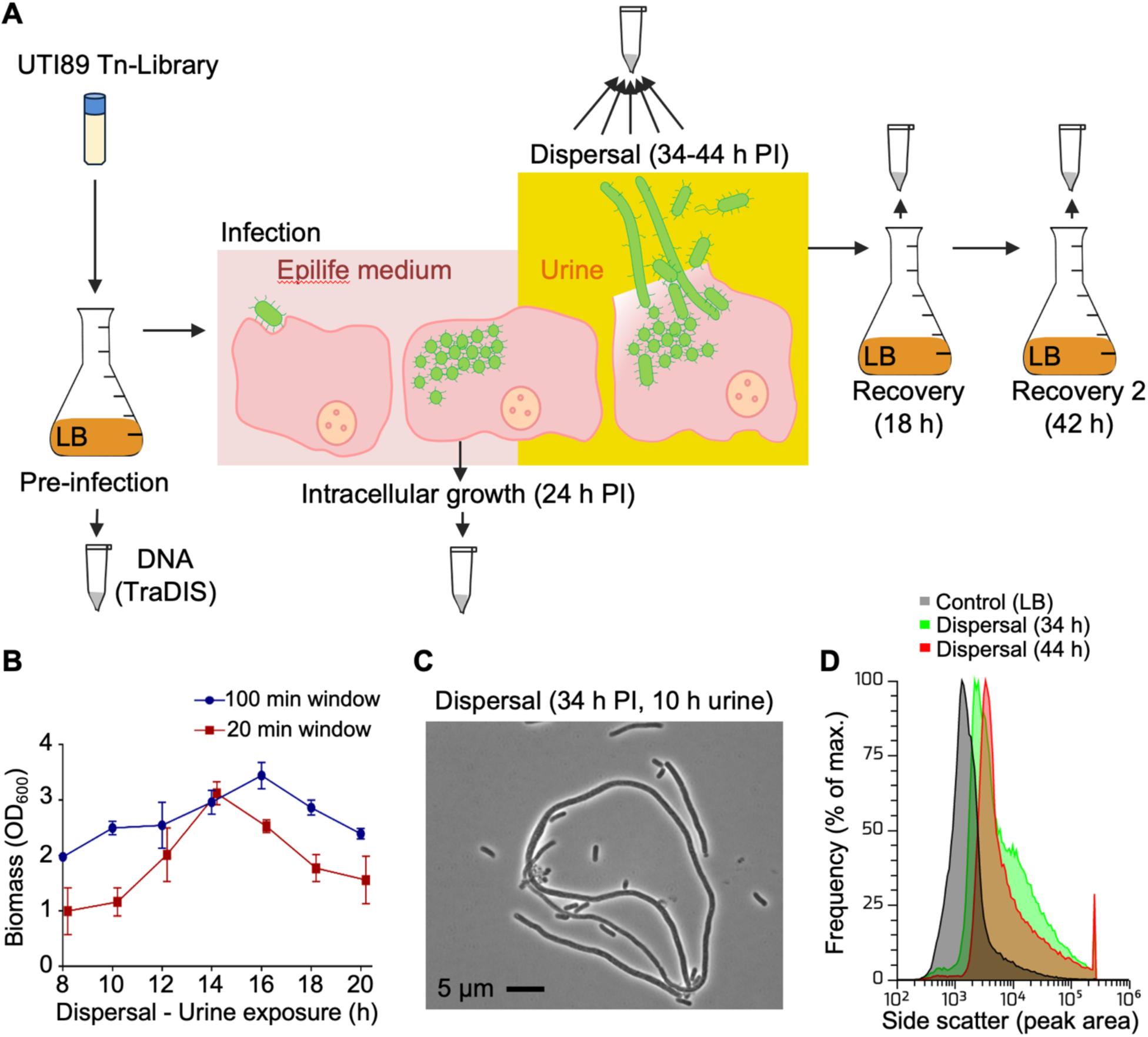
Cell culture model of UTI89 infection of BECs for TraDIS analysis. (A) The stages of infection and the pre-and post-infection samples that were subsequently analysed by TraDIS are represented by sample tubes. (B) Optical density measurements (OD_600nm_) of samples harvested in 20-or 100-min time windows during the urine exposure (dispersal) phase in the infection model. Error bars indicate the standard deviation. (C) Phase-contrast micrograph of a dispersion sample shows a mixture of rod-shaped and highly filamentous UPEC. (D) Flow cytometry frequency distributions of cell size (represented by the side-scatter peak area, arbitrary units), showing substantial populations of high-scatter filamentous cells during dispersal.

Samples from all the main infection stages (Figure 4A) were obtained in duplicate for TraDIS; **Supplementary Table 1B** shows read count statistics, and **Supplementary Table 1H** shows each gene’s Tn-specific read counts. Examples of the distribution of reads for selected loci are shown in **Figure 5A** and **5B**. These results suggested that certain mutants were selectively lost at different stages of infection in both replicates, as would be expected for genes important for the corresponding stage of infection. For some genes, we noted differences in the change in read counts over the course of infection between the replicate experiments (see Figure 5; blue and red bars represent data from separate replicates). For example, Tn-linked reads in the *hisF* gene appeared moderately reduced compared to its neighbouring genes in the IBC sample of one experiment, but they were almost completely missing in the other replicate (Figure 5A). We therefore chose to further investigate the *hisF* gene as an example in infection experiments with individual knockout strains (further below). We also applied the more stringent criteria (*i.e.*, *p*-value ≤ 0.05 and log_2_FC of ≤ -1.0, in both replicates) to improve the accuracy of gene function identification based on the results above.

**Figure 5.**
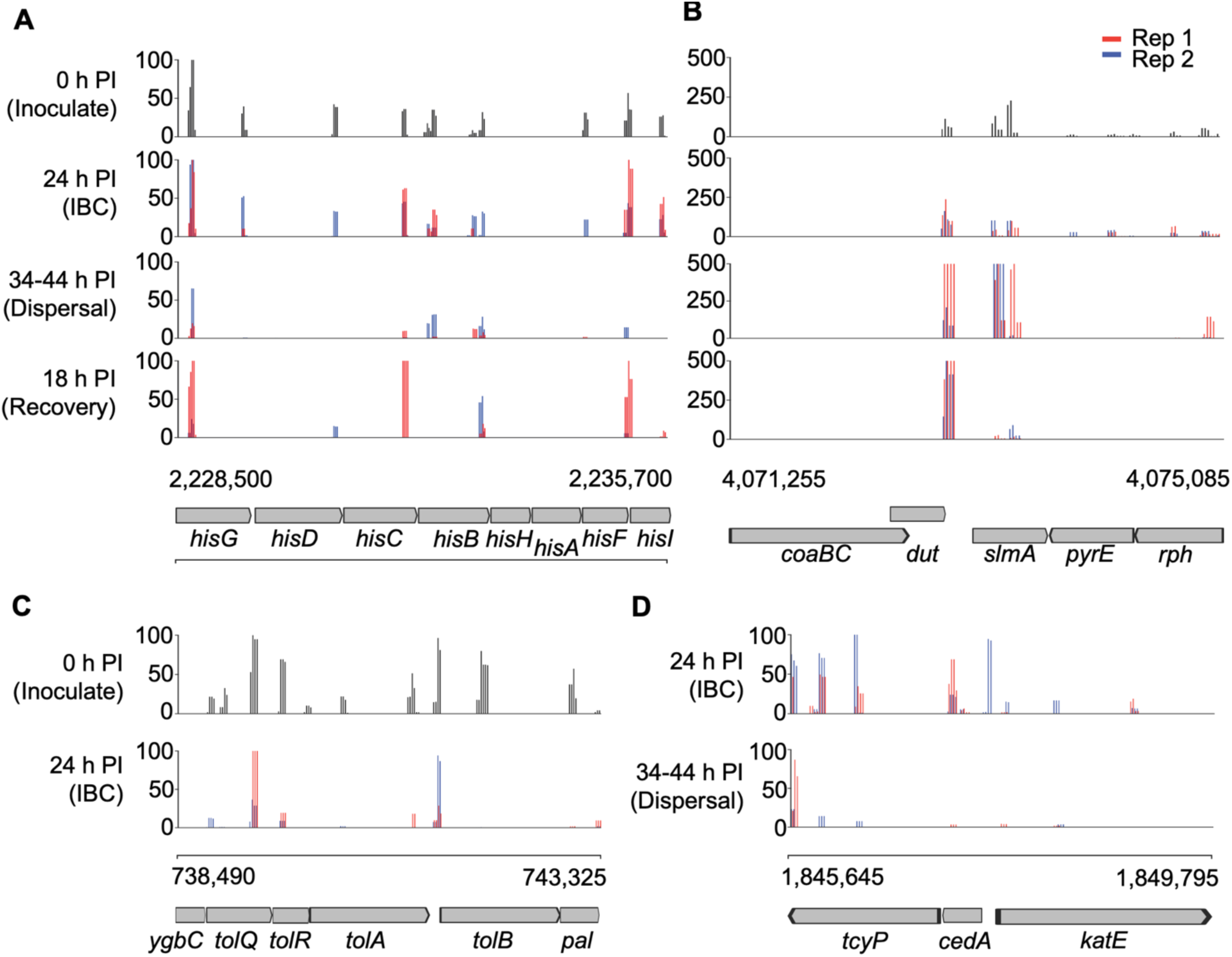
TraDIS read counts within selected UTI89 genomic regions at the indicated stages of bladder cell infection and recovery. (A-B) Mapped read counts in the indicated genomic regions in all stages of the infection and recovery process that were sampled. The two experimental replicates are shown with blue and red bars. (C) Mapped read counts from the Inoculate (top panel) and IBC (bottom panel) samples encompassing the indicated *tol* and *pal* genes. (D) Mapped read counts from the IBC (top panel) and Dispersal (bottom panel) samples encompassing genes from the *cedA* region. Each plot has a bin size of 50 bp. PI = post-infection.

### UTI89 genes implicated in intracellular growth in BECs (IBC stage)

To identify candidate genes required for intracellular infection, the TraDIS read counts for each gene obtained from the 24 h IBC samples were compared to those from the pre-infection sample. The full TraDIS data analyses for the IBC stage of infection are shown in **Supplementary Table 2A**. A total of 41 genes met the stringent criteria in both replicates (**Table 2**). Some of these genes are related to metabolism, precursor biosynthesis and cell size control (e.g., *galU*), amino-acid biosynthesis and transport (e.g., *aroK*), and iron acquisition (e.g., *iroE*) (**Table 2** and **Supplementary Table 2A**), which is consistent with a reliance on *de novo* biosynthesis of some essential metabolites, amino acids, and sugars during intracellular infection.

**Table 2.**
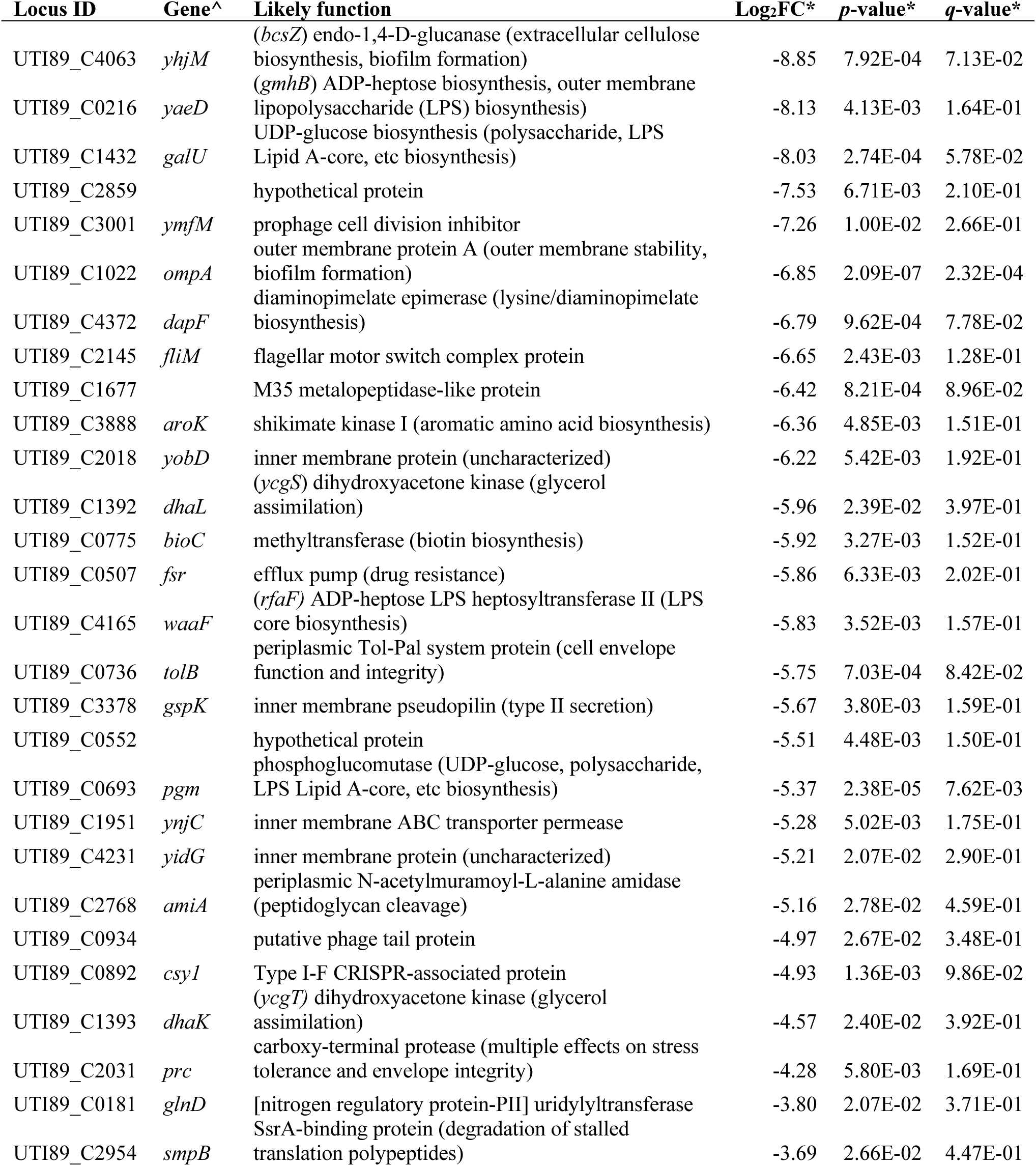
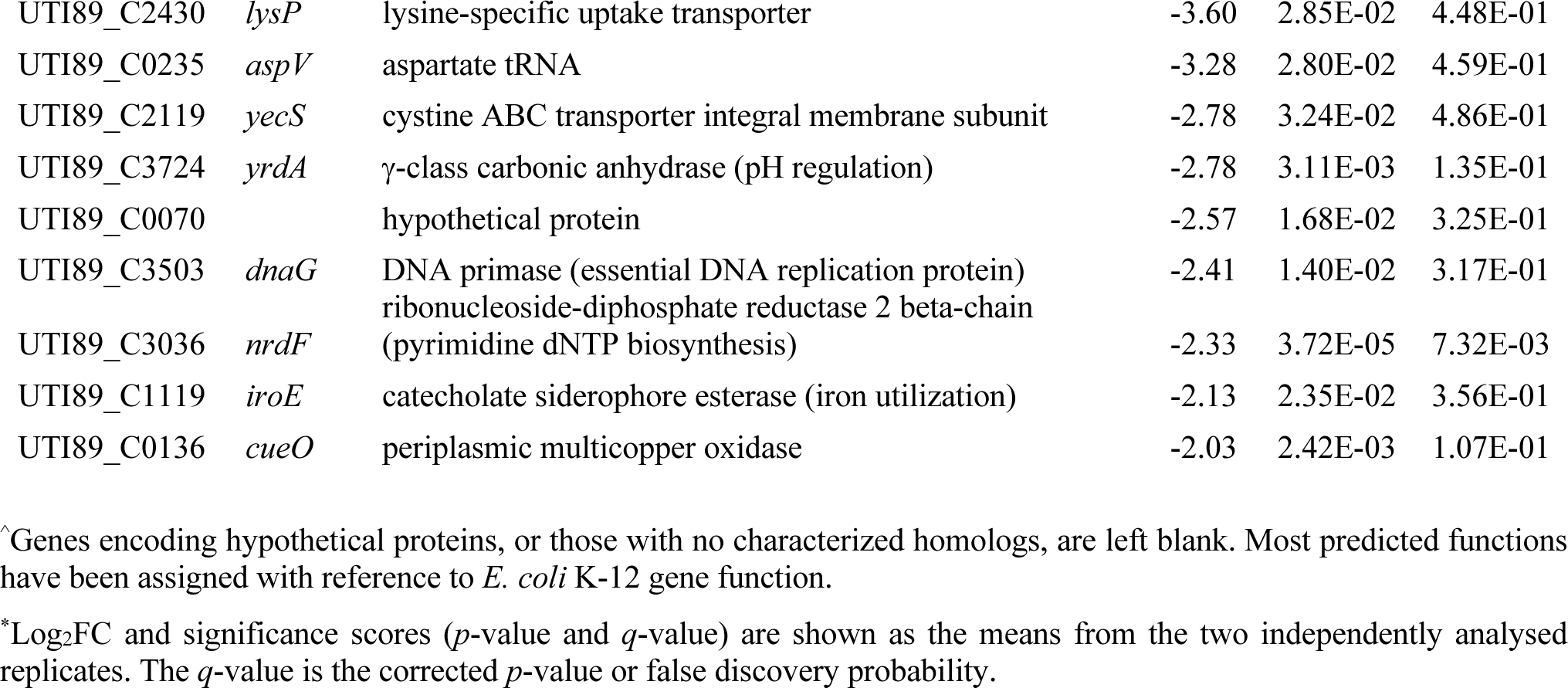
*E. coli* UTI89 candidate genes important for intracellular growth (IBC stage) compared to pre-infection LB culture.

Many of the TraDIS results above were consistent with certain mutants becoming selectively lost at specific stages of bladder cell infection. Interestingly, various macromolecule precursor and polysaccharide biosynthetic genes appeared to be important for both the intracellular stage of infection and for growth in M9-glycerol. We chose to independently test the importance of the *hisF* and *neuC* genes, which are important in M9-glycerol (Figure 3), for their potential role in intracellular stage (IBC) of infection, as implied by the TraDIS results albeit not in the high-stringency list (**Table 2**, **Supplementary Table 2A**). Infection experiments were carried out with WT UTI89 and ϕ..*hisF* and ϕ..*neuC* strains. After the IBC stage of infection, we observed a significant reduction in bacterial counts in both the ϕ..*hisF* and ϕ..*neuC* strains when compared to the WT (Figure 6), supporting the TraDIS results suggesting both the *hisF* and *neuC* genes are important for intracellular infection of BECs.

**Figure 6.**
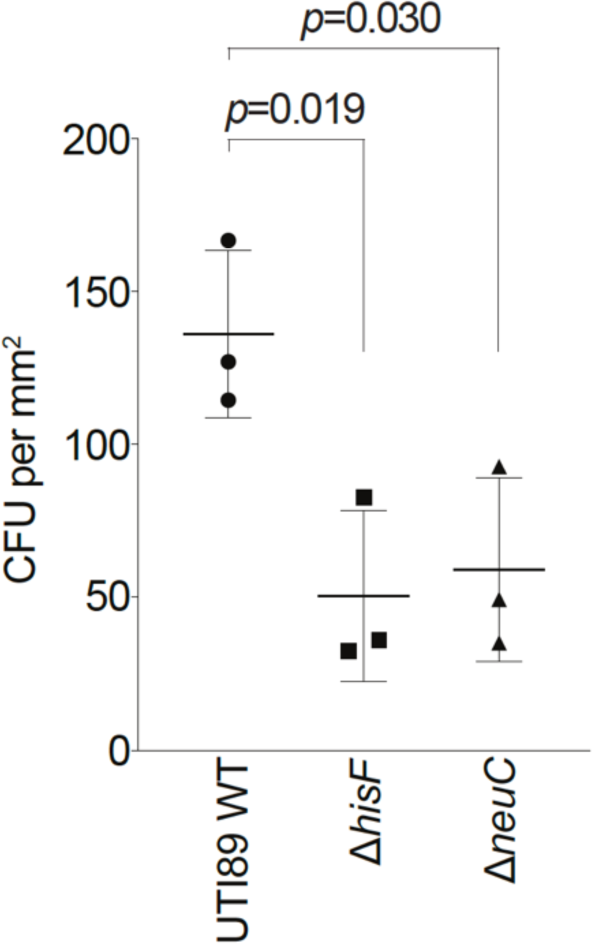
Deletion of *hisF* or *neuC* affects bladder cell infection. Gentamicin protected bacteria (intracellular) of the indicated strains were recovered at the IBC stage and colony forming units (CFU), per area of the infection surface, were counted. Error bars indicate the standard deviation (n = 3). Significance was calculated using a two-sided *t*-test.

### UTI89 genes implicated in UPEC dispersal from infection of BECs

When IBCs reach an advanced stage, conditions can trigger bacterial dispersal from host cells. UPEC may differentiate into highly filamentous or motile forms at this stage to aid dispersal [31]. To identify genes involved in progression to this poorly understood phase, we obtained bacteria released from the bladder cells by collecting the culture supernatant between 34-44 h post-infection (during exposure to human urine of density > ∼1.025 g/mL, which triggers dispersal). The resulting TraDIS data were analysed by comparison to the IBC data, to identify candidate genes specifically important for dispersal; the complete analysis output is given in **Supplementary Table 2B**, and a total of 148 genes met the stringent criteria in both replicates (**Supplementary Table 2C)**. Some of these genes have been implicated in the regulation of cell division (*cedA*, *ytfB*, *dedD* and *zwf,* and *ftsX*), and cell wall remodelling (*dacA*), along with other genes implicated in stress and signalling response pathways (**Supplementary Table 2C)**. Notably, while the *waaL* gene involved in LPS biosynthesis also appeared in this list, *waaL* mutants diminished rapidly over time in this library even in LB (Figure 1, **Supplementary Table 1F**).

### UPEC genes implicated in bacterial recovery from infection of BECs

Upon dispersal, filamentous UPEC can revert to rod cells through unknown mechanisms [25, 32]. This is thought to be necessary for long-term bacterial survival and for regaining invasive capacity to initiate a new cycle of infection. To search for genes important for bacterial recovery after dispersal, bacteria harvested from the dispersal phase were collected and then resuspended in LB and allowed to grow for 24 h (recovery sample). A sample of this culture was then grown for a further 24 h in liquid LB media (48 h recovery sample). TraDIS data obtained from these recovery samples were then separately compared to the dispersal TraDIS data, to identify any mutants that had a significantly reduced ability to recover from infection (**Supplementary Table 2D** and **2E**). A total of 283 and 313 genes met the stringent criteria (*p*-value cut-off of ≤ 0.05, log_2_FC of ≤ -1.0, in both replicates) at the 24 h and 48 h recovery samples, respectively (**Supplementary Tables 2F** and **2G**). A total of 233 genes were found in common between the 24 h and 48 h recovery samples, and 40 genes were found specific to the 24 h recovery sample, which included some implicated in the regulation of cell division, envelope stress response and peptidoglycan remodelling, including *yfiR, yibP* (*envC*) and *ampH* (**Supplementary Table 2F**), consistent with likely requirements for bacterial recovery from envelope stresses experienced during intracellular infection [32]. For example, YfiR is known to promote UPEC survival in the mouse model of UTI [33], and is thought to be a negative regulator of YfiN (DcgN), which inhibits cell division in response to cell envelope stress [34]. Further work will be needed to ascertain the potential specific role of many of these genes in the infection cycle.

### Testing genes implicated in bacterial dispersal and recovery from bladder cell infection

To facilitate the further direct testing of the role of genes implicated in infection, we next established a direct competition assay of infection, where wild-type UTI89 and a mutant were differentially labelled with cytoplasmic fluorescent proteins (GFP or mCherry) and then cultures equally mixed before infection; the proportions of the two strains may be determined at pre- and post-infection stages by fluorescence microscopy. The direct comparison of WT and mutant in the one infection was expected to reduce experimental variance and improve throughput. We applied this method to two genes of interest identified in the dispersal and recovery stage screens, *dedD* and *slmA* respectively (**Supplementary Table 2C and 2E**), which have roles in regulating cell division. As may be seen in Figure 7, the UTI89 11.*dedD* strain grew well against wild-type UTI89 in LB co-culture over the same period as used in the infection model (Figure 7C) but was strongly out-competed in samples taken in the dispersal stage of infection (Figure 7B), consistent with TraDIS screen. In separate infection experiments, the 11.*dedD* strain produced filaments visually indistinguishable from wild type on dispersal (**Supplementary Figure 5**), further suggesting that *dedD* plays a role in maintaining growth rate or survival during dispersal rather than directly regulating filamentation.

**Figure 7.**
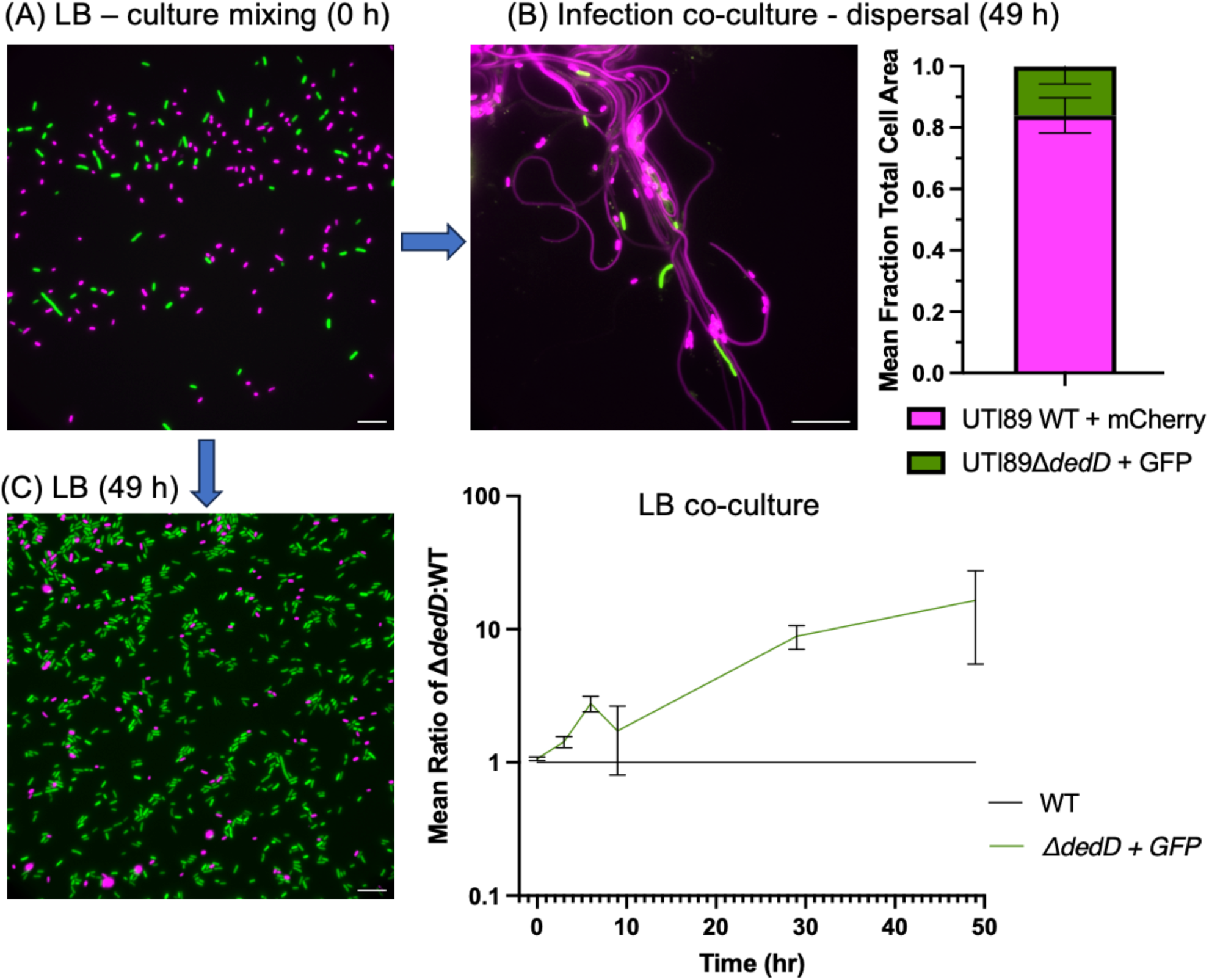
Deletion of *dedD* affects bacterial growth during infection of BECs. (A) Fluorescence composite image of mixed UTI89 WT (mCherry - magenta) and UTI89.Δ*dedD* (GFP - green) strains (pre-infection, 0 h). (B) The relative biomass (total cell area) was measured after dispersal (20 h urine treatment) from infection (49 h) (n = 3). (C) LB co-culture assay, indicating a growth advantage of UTI89.Δ*dedD* (GFP) compared to WT (mCherry); the example image at 49 h shows UTI89.Δ*dedD* (GFP) dominant. Cell area data were normalized at each timepoint using a WT (mCherry) versus WT (GFP) control co-culture, normalized to a ratio of 1 (n = 3). Cultures were passaged at 9 h and 29 h (1:1000 dilution into fresh LB).

SlmA helps prevent cell division occurring over the DNA during the *E. coli* cell cycle, although a single *slmA* deletion shows a weak phenotype [35]. Since DNA forms an irregular extended axial filament within UPEC filaments [25], we hypothesised *slmA* could be especially important for proper division site positioning between DNA as nucleoids condense during filament reversal. However, initial results comparing UTI89 wild-type and 11.*slmA* during infection recovery were inconclusive and showed no obvious defect during the infection-recovery phase. We therefore did time-course imaging to visualize the division of filaments and DNA localization during recovery in LB. This also showed no major defects of UTI89.11.*slmA* in DNA condensation into distinct nucleoids or filament division (**Supplementary Figure 6, Supplementary Movie 1**). Further studies will be required to investigate any possible alternative or relatively minor roles of *slmA* at this stage, and any of the other potential stage-specific gene functions that have been predicted based on the TraDIS screens presented here.

## DISCUSSION

To screen for bacterial genes important for infection of host cells in culture, we have developed a modified TraDIS protocol that uses regular Illumina sequencing and is amenable to such analyses where culture-scale and DNA yield may be limited. We applied this method to a transposon insertion mutant library of *E. coli* UTI89 to identify genes important during growth in M9-glycerol minimal medium, and during the main stages of infection of cultured bladder epithelial cells by comparing TraDIS data obtained from each stage of infection to the preceding one. This approach should facilitate comprehensive genome-wide analyses of gene function during stages of other microbial infections or complex developmental pathways.

In characterizing the UTI89 transposon insertion library, we identified 196 genes meeting a significance threshold that tolerated transposon insertions during the initial library generation on LB-agar but were important for ongoing growth (5 overnight passages) in LB liquid culture. Amongst these were several genes involved in biosynthesis of the lipopolysaccharide (LPS) core of the *E. coli* outer membrane, such as *waaY*, *waaL* and *waaV* (or *rfa* genes), likely reflecting the general importance of outer-membrane stability to cell survival and ongoing growth in culture [36]. Some of these genes were also identified in the TraDIS screen for survival during infection of bladder cells, when compared to a pre-infection sample of the library. This might reflect their general importance over time in culture, rather than an infection-specific role, and highlights the value of growing the Tn-library for the same number of generations in each condition for comparison in Tn-seq studies. In infection studies, where this may not be practical or possible to do accurately, comparing the screen results to those of batch cultures over time would therefore help assess the likelihood that identified genes have infection-specific roles.

We also identified 16 genes where Tn-insertions appeared to improve the fitness of the bacteria over time in LB, some of which may reflect a false discovery rate and an anticipated limitation due to the scale of the Tn-library and infection model that were feasible in our study. Yet others might reflect a genuine enhancement of growth due to gene inactivation, which potentially includes some transcriptional repressors of genes involved in nutrient utilization (e.g., *cytR*, Figure 1E), like those seen in the original *Salmonella* TraDIS study [10]. We speculate that these types of genes are dispensable in rich medium, where they could limit maximal nutrient utilization and growth, but would be beneficial for regulating nutrient use in natural environments.

Natural environments like a host cell cytoplasm require certain *de novo* biosynthesis and nutrient utilization functions of infecting bacteria, including during UTI [5, 37–39]. Our screens for UTI89 genes required for transition to and growth in M9-glycerol minimal medium and for intracellular growth during infection expand on this view and showed common types of gene requirements. Many of the strongly identified genes in M9-glycerol and intracellular conditions are implicated in macromolecular precursor biosynthesis, such as amino acids, purines, and pyrimidines, or in assimilation of nutrients that are consistent with substantial nutrient limitations for bacteria in the host intracellular environment (**Tables 1 and 2**). Independent gene deletion studies in both UTI89 and K-12 strains based on the UTI89 screen results verified the relevance of our genetic screens in the identification of gene functional requirements in M9-glycerol medium and showed a clear congruence between the specific genetic requirements of both strains, as expected. We verified that the histidine biosynthesis gene, *hisF*, was essential for growth in M9-glycerol and important for intracellular infection of bladder cells but was completely dispensable in the rich LB medium. This and other bacterial biosynthetic genes strongly identified in our study during infection show the diverse requirements for macromolecular precursor biosynthesis during intracellular infection by UPEC.

Genes involved in cell envelope stability and stress tolerance were also found in both M9-glycerol and intracellular infection screens (**Tables 1 and 2**). These included outer membrane stabilization factors, such as the Tol-Pal complex, which have been broadly implicated in virulence. Of particular apparent importance to intracellular infection by UTI89 were genes involved in a range of cell surface polysaccharide biosynthesis, cell envelope stress tolerance and biofilm-related functions (**Table 2**). Interestingly, we found that *neuC*, which is required for biosynthesis of sialic acid as a precursor for extracellular K1 capsule polysaccharide in some pathogens [12, 40], is important for UTI89 intracellular infection of human bladder cells. This is consistent with a previous report of K1 capsule involvement in IBC formation in the mouse model of cystitis [41]. The role of capsule in UPEC has otherwise been linked to resistance to phagocytosis, immune complement, and survival in blood [12, 42–44]. The exact role of *neuC* that improves growth in M9-glycerol is uncertain but, given the importance of other envelope-stability and stress related genes in our data (e.g., *tol-pal*, *rpoS*), this might reflect potential stabilisation of the outer cell envelope by the capsule during several of these stress conditions.

The requirement for capsule in various conditions appears to be strain dependent, as, for example, *E. coli* K-12 does not encode the K1 sialic acid capsule but grows well in M9-glycerol (e.g., Figure 3). Both K-12 and UTI89 have another UDP-*N*-acetylglucosamine 2-epimerase distantly related to NeuC called WecB (RffE). However, these proteins are involved in different surface polysaccharide biosynthesis pathways and the genes are not complementary [12, 40, 45]. This is consistent with our observation that *neuC* is important for growth in M9-glycerol, despite the presence of *wecB* in UTI89. Overall, these results suggest that K1 capsular polysaccharide has multiple functions, not all of which may be needed in different strain contexts, consistent with the evolving view that gene function is highly strain and context dependent [29].

Our study also searched for genes that function in the latter stages of cellular infection, where bacteria erupt from host cells and disperse into the extracellular environment in differentiated forms, such as highly prevalent filamentous bacteria and motile rods [4]. Infection-related filamentation (IRF) occurs by UPEC cell growth without division in response to multiple conditions encountered during UTI, including urine-specific factors [3, 25, 46]. IRF is a likely stress response that may aid in bacterial dispersal [31]. The subsequent recovery steps, including filament reversal (division to rods), are thought to be important to allow long-term survival and potential re-infection of other host cells [4, 32]. Our observations of envelope integrity and stress response gene requirements for survival of intracellular UPEC (**Table 2**), and that some bacteria lyse upon dispersal [32], suggest that envelope stress is a major challenge to bacterial survival during UTI. Consistent with this, genes implicated in cell division and envelope regulation were observed in our dispersal and recovery stage screens results.

One relatively strongly predicted gene in our dispersal screen was *ytfB*. This gene has roles in both cell division (in K-12) and binding to host cell surface glycans (in UTI89) [47, 48]. A substantial effect of *ytfB* on cell division only became evident when 11.*ytfB* was combined with 11.*dedD* [48], which is thought to play a role in peptidoglycan stabilization during cell division [49]. Interestingly, we also found that *dedD* has an important role during dispersal from host cells, which was verified by a fluorescence-based competitive infection assay (Figure 7). Since YtfB and DedD both have extracytoplasmic glycan-binding capacities [47–49], we speculate that these proteins play roles in stabilizing peptidoglycan and cell division or the cell envelope during envelope stress or conditions experienced by UPEC in the latter stages of the infection cycle.

Many genes we identified with potential roles in the dispersal and recovery stages of infection have diverse predicted functions or are currently poorly characterized and warrant future investigation. Additional studies on genes identified in the screens here are important before conclusions should be drawn about their roles because we observed a moderate false-discovery rate due to the necessarily limited library size and infection scales that were feasible in our study. The genes identified here nevertheless represent a spectrum of potential targets for therapeutic intervention in UTI.

## METHODS

### Bacterial strains and standard growth conditions

*E. coli* BW25113 wild-type and Keio knock-out strains harbouring individual mutants of *hisF*, *argC*, *bioF*, *yggB*, *ykgC*, *sgbE* and *pdhR* were obtained from the *E. coli* Genetic Stock Center (Yale University, USA). Uropathogenic *E. coli* UTI89 [2] knock-out mutants were constructed using λ-Red recombination (29). Primers containing flanks for red-recombination and 3’ ends for amplifying the kanamycin cassette from pKD4 are given in **Supplementary Table 1A**. Mutants were selected using LB agar supplemented with kanamycin (50 μg/mL) and confirmed using allele-specific PCR with external and internal primers to the kanamycin resistance gene (primer K1, **Supplementary Table 1A**).

Strains were routinely cultured at 37°C on solid or in liquid LB (including 1 % (w/v) NaCl), unless otherwise stated. M9 minimal medium was also used where indicated, including 1 % (v/v) glycerol as the carbon source, and a trace elements solution (28). For culturing the UTI89 transposon mutant library, kanamycin (50 μg/mL) was included.

### Construction of a mini-Tn5 transposon mutant library in UTI89

*E. coli* UTI89 was prepared for electroporation as previously described (30). A volume of 50 μL of electro-competent UTI89 was mixed with 1 μL of EZ-Tn5 <KAN-2> Tn5 transposome mix (Epicenter Biotechnologies) and placed on ice for 5 min in a 2 mm electroporation cuvette. The cells were then electroporated using a BioRad GenePulser, with settings of 2.5 kV, 25 μF and 200 Ω. We then added 950 μL of SOC medium (30) to the cells, and then eleven replicate electroporated samples were then pooled and incubated at 37 °C for 1 h. The culture was then concentrated and to 1 mL by centrifugation and resuspension, and then spread onto LB-agar supplemented with kanamycin (50 μg/mL). The total number of mutant colonies was measured by plate counting of 1/100 and 1/500 dilutions of the main culture (total of 97,500; SEM = 882, n = 3). All plates were incubated for ∼18 h at 37°C. Colonies (lawn) were then resuspended with 1 mL of LB supplemented with kanamycin (50 μg/mL) and 15 % (v/v) glycerol, then pooled and mixed. This library was stored in 1 mL aliquots at -80 °C.

### Transposon-directed insertion-site sequencing (TraDIS)

We used Tn5-transposasome mediated DNA fragmentation coupled to linker attachment (tagmentation) and dual-end transposon insertion-site PCR (**Supplementary Figure 1A**), to significantly reduce the amount of input DNA required and eliminate the need for custom sequencing primers and dark cycles during sequencing with the original TraDIS method [10]. Genomic DNA (gDNA, 5 ng) from the transposon mutant library (grown as indicated above in LB, M9-glycerol or at distinct phases of infection) was subjected to tagmentation using the Nextera DNA Library Prep Kit (Illumina). The DNA was mixed with 10 μL of 2x TD Buffer and 5 μL of Tagment DNA Enzyme TDE1 (diluted 1/50 in 0.5x TE buffer and 50 % (v/v) glycerol) and made up to a final volume of 25 µL, and then incubated at 55°C for 5 min. A volume of 5 μL of 0.2 % (w/v) SDS was then mixed with the sample to stop the tagmentation reaction and the sample was incubated for 5 min at room temperature.

To amplify the transposon-genomic DNA junctions from both transposon ends separately, each PCR combined a transposon-specific outward-directed primer with a primer that anneals to one of the end-adapter sequences (**Supplementary Figure 1B**). Both primers include 5’ tag sequences that make them suitable for downstream sequencing **(Supplementary Table 1A)**, as described below. The transposon-specific primer was designed to include (from 5’ to 3’): (i) the “P5” or “P7” flow-cell annealing sequence (Illumina), (ii) a standard “read 1” or “read 2” sequencing primer binding site, and (iii) a ∼20 bp transposon-specific annealing sequence, designed to amplify ∼40–50 bp of the corresponding transposon end followed by its adjacent genomic sequence. Two such transposon-specific primers were designed—one for each end of the transposon; one end was represented by the P7/read 1 tag, and the other by the P5/read 2 tag (**Supplementary Figure 1B**). The second primer in each PCR reaction follows the standard design for Illumina sequencing, containing (from 5’ to 3’): (i) the alternate P5 or P7 adapter sequence for binding to the Illumina sequencing flow cell, (ii) a unique barcode sequence for later sample identification (de-multiplexing), and (iii) a sequence that anneals to the tagmentation adapter on the opposing end of the fragment, which then serves as the read primer binding site during sequencing. The dual Tn-end sequencing design ensured that only ∼50% of reads contained a common Tn-end sequence during the early cycles of each paired-end read, and the remaining reads comprised genomic sequence reads without Tn-end sequences (**Supplementary Figure 1C**).

The PCR reactions were carried out using the KAPA HiFi Library Amplification Kit (KAPA Biosciences). Primers (0.2 μM each) were added directly to the tagmentation reaction along with the KAPA HiFi HotStart ReadyMix. As a control for non-specific amplification, identical PCR reactions containing a single primer were also performed and did not generate significant amplified product. An initial extension step was performed at 72°C for 3 min. This was followed by a denaturation step for at 98°C for 30 sec, followed by 28 cycles of: 98°C for 15 sec, 55°C for 30 sec, and 72°C for 30 sec. A final extension of 72°C 5 min was performed.

The purified PCR products were subjected to standard Illumina paired-end sequencing (**Supplementary Figure 1C**). Multiplex sequencing with samples from both Tn ends together provided the necessary read diversity during Illumina sequencing, avoiding the need for custom dark cycles. The amplified DNA samples were quantified with an Agilent Bioanalyzer and pooled to equimolar concentrations and underwent SPRI-select magnetic bead clean-up (Beckman-Coulter) at 0.9-0.5x left and right ratios, respectively (to select DNA of ∼200-800 bp). The pooled sequencing libraries were quantified on an Agilent Bioanalyzer. PhiX genomic adapter-ligated DNA control library (Illumina) was included at 5% of the total DNA. DNA was then denatured and 10 pM was sequenced on a V3 paired end (PE) Illumina flow cell using an MiSeq sequencer for 300 cycles (2 x 150 nt paired-end reads) for M9-glycerol or on an Illumina HiSeq 2500 sequencer for 250 cycles for the infection samples, according to the manufacturer’s instructions.

### Analysis of TraDIS data

Sequence reads from the multiplex FASTQ sequencing data files were separated into one file for each sample, according to their unique index barcodes. The data underwent quality control filtering using FastQC v0.11.2 (Simon Andrew, 2010, http://www.bioinformatics.babraham.ac.uk/projects/fastqc/). Sequence reads containing a match to the designated i5 and i7 Tn-ends were isolated, allowing up to 3 mismatches (**Supplementary Table 1B**), trimmed of the Tn-end sequence, and then mapped to the genome of UTI89 (chromosome and pUTI89) using the SMALT short read mapper, allowing no mismatches, as described previously [50].

The number of mapped reads in bins along the genome sequence was determined with deepTools2 [51]. The number of Tn-insertion sites for each gene and the comparative analyses between two conditions were done using Bio-TraDIS [50]; the two read files for both Tn-ends were input separately for each biological sample. Another analysis was then conducted for the second culture replicate. Genes that contained ≤ 5 reads were removed from the comparison. Genes meeting the defined statistical thresholds (*i.e.*, *p*-value ≤ 0.05 and log_2_FC of ≤ -1.0) in both replicate cultures were identified (**Table 1 and 2**); the Log_2_FC and significance scores (*p*-value and *q*-value—the false discovery rate) were given as the means from the two independently analysed replicates.

### Culturing the UTI89 library in LB and M9-glycerol for TraDIS

To carry out five sequential passages of the UTI89 mutant library in LB medium, an aliquot of the UTI89 transposon mutant library was first thawed at room temperature, and a sample was taken for DNA extraction and purification using an Isolate II Genomic DNA purification kit (Bioline). Also, 50 µL of the thawed library aliquot was used to inoculate 20 mL of LB and this culture was incubated for 24 h at 37°C with 180 rpm shaking (passage 1). A volume of 50 µL of the passage 1 culture was then used to inoculate 20 mL of LB and incubated the same way for a further 24 h (passage 2). This was sequentially repeated for a total of five consecutive passages. After each passage, 10 mL of each culture was harvested for genomic DNA extraction and purification.

To compare the growth and survival of the *E. coli* UTI89 transposon-insertion mutants in M9-glycerol and LB media, an aliquot of the mutant library was thawed at room temperature and diluted in LB (to 150,000 cfu per mL), which was then incubated at 37°C for 1.5 h. One mL from the LB starter culture was then added to 9 mL LB, and another 1 mL of the starter culture was added to 9 mL M9-glycerol. These cultures were incubated for 24 h at 37°C with 200 rpm shaking. A volume of 1 mL from each culture was then used to inoculate 10 mL of the corresponding fresh medium and grown for another 24 h at 37 °C with 200 rpm shaking. Finally, 10 mL from this culture was then used to inoculate a final volume of 90 mL of fresh medium and grown for 24 h at 37°C with 200 rpm shaking. Samples were harvested for genomic DNA extraction and TraDIS. This sequential dilution from LB into M9-glycerol supported the transition between media and supplied sufficient auxotrophic requirements for UTI89 growth [52].

### Culture of individual deletion mutants in M9-glycerol and LB

The wild-type and knock-out mutant strains were streak-plated onto LB agar supplemented with kanamycin (50 μg/mL) and incubated at 37°C for ∼16 h. Single colonies were used to inoculate 5 mL of pre-warmed liquid LB (supplemented with kanamycin, where appropriate) and the cultures were incubated at 37°C with shaking for ∼16 h. A volume of 500 μL was then used to inoculate 5 mL LB or M9-glycerol medium and incubated at 37°C with shaking for ∼16 h. Two hundred microliter volumes of fresh LB or M9-glycerol medium, respectively, were placed into each well of a 96-well sterile microtiter plate, and then inoculated with the starter cultures with the corresponding media, to give a starting OD_600 nm_ of 0.015. The plate was incubated at 37°C for 24 h with shaking and analyzed with a microtiter plate spectrophotometer (BioTek), measuring the OD_600 nm_ at 30 min intervals. Sterile medium wells were used as the blank reference. The growth curves were obtained by averaging technical replicates within two independently performed microtiter plate experiments. The DMFit (DM: Dynamic Modelling, version 3.5) growth curve modelling software (34) was used to obtain values for the lag time, growth rate (μ) and maximum OD.

### Infection of bladder epithelial cells with the UTI89 library for TraDIS

Human bladder cell line PD07i was cultured at 37°C with 5 % CO_2_ to confluency [3, 25]. Bladder cells were trypsinized, concentrated to 4 x 10^5^ /mL, and then 10 mL was dispensed into each sterile tissue culture dish (BD Falcon, 100 mm diameter x 20 mm depth) and incubated at 37°C with 5 % CO_2_ for 24 h. This method below is described at a scale of 1 dish, but the infection can be readily scaled to include multiple dishes.

#### IBC phase of infection

An aliquot of the UTI89 transposon mutant library was thawed at room temperature and 100 μL was used to inoculate 10 mL of liquid LB media and grown statically overnight at 37°C. The bacterial culture was then diluted down to OD_600 nm_ = 1.0 in PBS and 1 mL of the dilution was used to inoculate each bladder cell culture dish. The culture dishes were then centrifuged in a swinging-bucket rotor for 5 min at 500 g and left to incubate statically for 2 h at 37°C with 5 % CO_2_.

The supernatant was then discarded, and 10 mL pre-warmed EpiLife medium (Thermo-Fisher) supplemented with human keratinocyte growth supplement (HKGS) and gentamicin (100 μg/mL). Culture dishes were then incubated statically for 22 h at 37°C with 5 % CO_2_. Where necessary, culture dishes were imaged in phase-contrast and GFP fluorescence at 18 and 22 h using a Nikon Ti inverted microscope. After 22 h of static growth the supernatant was removed, and the culture gently washed 4 times with PBS. 1 mL of lysis solution (0.5 % trypsin-EDTA and 0.1 % triton X-100) was added and incubated for 10 min at 37°C. The entire supernatant was then collected and spun at 4,600 rpm for 10 min. The supernatant was removed, and the cell pellet underwent gDNA extraction and purification.

#### Dispersal phase of infection

To induce the IBC-dispersal stage of infection and mimic the main lifecycle stages of UPEC in an *in vivo* UTI, the infected bladder cells were exposed to human urine from a mildly dehydrated male donor (density > 1.025 g/mL). Urine was centrifuged at 4,600 rpm for 10 min and the supernatant was then filter sterilised (with a 0.2 μm filter). Filtered urine samples were then pooled (a density of 1.026 g/mL was measured for the experiments described here), and stored in 40 mL aliquots at -20°C. Each aliquot was thawed at 37°C before use. Following 22 h of intracellular infection, as described above, the Epilife medium supernatant was removed, and the dishes were gently washed 4 times with PBS. 10 mL of the urine (supplemented with kanamycin 50 μg/mL, when infecting with the transposon library only) was added and the culture dish left to incubate at 37°C with 5 % CO_2_ and gentle orbital shaking (∼50 rpm to resuspend bacteria released from the bladder cells). During incubation, the overlay urine was periodically withdrawn for analysis and immediately exchanged for a new sample of pre-warmed urine, after incubation for 20 min windows over 10 h. The short windows of collection aimed to minimise reversal and recovery of the mutants in the urine (Figure 4C), so that genes required for these subsequent stages would not be falsely detected in the dispersal sample. The collected samples were placed on ice, centrifuged at 4,600 rpm and the cell pellets frozen. The next day, the cell pellets were resuspended and pooled for gDNA extraction and purification.

#### Bacterial recovery from infection

From the 10 h time point of urine exposure (34 h PI), urine was collected in 2 h collection windows (pre-warmed urine was re-applied after each collection), centrifuged, and stored on ice. Once all samples were collected, the cell pellets were resuspended in 10 mL of warm liquid LB media. Cultures were pooled and then incubated for 18 h overnight at 37°C with 180 rpm shaking. 20 mL of this culture was then centrifuged at 4,600 rpm for 10 min and the cell pellet underwent gDNA extraction and purification. In addition, a further 1/500 dilution into liquid LB media of the 18 h recovered culture was done, and the culture incubated for a further 24 h with 180 rpm shaking. This aimed to add an additional selection step for identifying mutants that had recovered. Following the extra 24 h growth, the culture was centrifuged at 4,600 rpm for 10 min and the cell pellet underwent gDNA extraction and purification.

### Infection assay of hisF and neuC deletion strains

Human bladder cell line PD07i was cultured at 37°C with 5% CO_2_ to confluency as previously described and counted using a Coulter Counter (Beckman) to a minimum of 1×10^5^ cells/mL. 0.5% Trypsin-EDTA mixture and defined trypsin inhibitor (Gibco) was used to release bladder cells from base of flasks. Bladder cells were concentrated to 4 x 10^5^ /mL and 1 mL was dispensed into a 6-well sterile culture dish (BD Falcon; 35 x 18 mm^2^) and incubated at 37°C with 5% CO_2_ for 24 h. Bacterial cultures for UTI89 WT, 11.*hisF* and 11.*neuC* strains were grown overnight statically and then diluted down to an OD_600 nm_ of 1.0. This was then used to inoculate the bladder cells by adding 0.1 μL/mm^2^ of bacteria to give a multiplicity of infection (MOI) of 100. The plates were then spun down for 5 min at 500 g and incubated statically for 2 h at 37°C with 5% CO_2_. Following this, the supernatant was discarded, and the bladder cells were resuspended in EpiLife supplemented with gentamicin (100 μg/mL) supplemented and HKGS. Cultures were then incubated statically at 37°C for 22 h. Following this, culture dishes were washed gently with PBS 4 times and lysis solution (0.5% trypsin-EDTA and 0.1% triton X-100) was added and incubated for 10 min at 37°C. The entire supernatant was then collected and plated onto LB-agar petri plates and left to incubate at 37°C for 18 h. Colonies were then counted from a 1/100-dilution series. This experiment and the results presented in Figure 6 was performed using 3 technical replicates.

### Growth competition (co-culture) assays

Wild-type UTI89 and each gene deletion mutant strain were transformed with pGI5 (GFP) or pGI6 (mCherry) [3, 53]. A single colony of each strain was used to inoculate LB medium, supplemented with 100 μg/mL spectinomycin, and the cultures were incubated at 37 °C. Each culture was then diluted to an OD_600 nm_ of 0.2 and mixed equally. This starting mixture was imaged by fluorescence microscopy (0 h sample).

For co-infection assays, a sample of an equally mixed bacterial culture was used to infect confluent PD07i bladder epithelial cells in flow chambers, as described [32]. In the 11.*dedD* experiment, we omitted the non-essential flushing step (100 μl.min^−1^) at the start of infection. For the reference LB co-culture assays, we passaged and sampled the co-culture at times corresponding to the main stages of the infection model. The mixed bacteria were diluted in LB (to OD_600 nm_ = 0.01) and then incubated at 37 °C with shaking at 200 rpm and sampled at 3, 6, and 9 h for microscopy. At 9 h, the culture was diluted 1:1000 and, after a further 20 h of growth (i.e., 29 h total), sampling for microscopy and culture dilution (1:1000) were carried out again. The diluted culture was incubated as above for a further 20 h before for imaging (49 h sample). The total area of cells of each strain in fluorescence micrographs was used as a measurement of the relative biomass of each strain. At each timepoint, the data for a control co-culture (WT-mCherry versus WT-GFP) were normalised to 50% each (i.e., a ratio of 1).

This normalization factor was then applied to the WT-mCh versus Δ*dedD*-GFP co-culture sample for each timepoint.

### Microscopy

Bacterial samples (1-3 μL) were placed onto 1-1.2 % (w/v) agarose pads (in PBS or LB) on a glass slide and a cover slip was placed on top. Phase-contrast or fluorescence imaging was performed with a Nikon Ti2-E microscope with a 100X 1.4 NA objective, and GFP or mCherry filter sets where appropriate. All image analysis was conducted in FIJI to determine the total fluorescence area of cells of each strain, representing biomass. Statistical analyses and data representation were done using GraphPad Prism 9.2.0.

## SUPPLEMENTARY DATA

Supplementary data figures (6) and tables (2) files are available with this article online. Nucleotide sequence data has been deposited under NCBI BioProject ID PRJNA1047296.

**Supplementary Movie 1**. **Deletion of *slmA* in UTI89 does not substantially affect filament reversal using recovery from infection.** UTI89 WT and 11.*slmA* strains were sampled during the dispersal stage from infections and placed on LB-agar pads for *in situ* visualization of recovery at 37°C by brightfield microscopy. Both strains show similar rates of filament division and growth.

## ACKNOWLEDGMENTS

The authors thank Hannah Brown for helping apply image analysis techniques. Part of this work was carried out during the award of an ARC Future Fellowship to IGD (FT160100010).

---

**Supplementary Figure 1.**
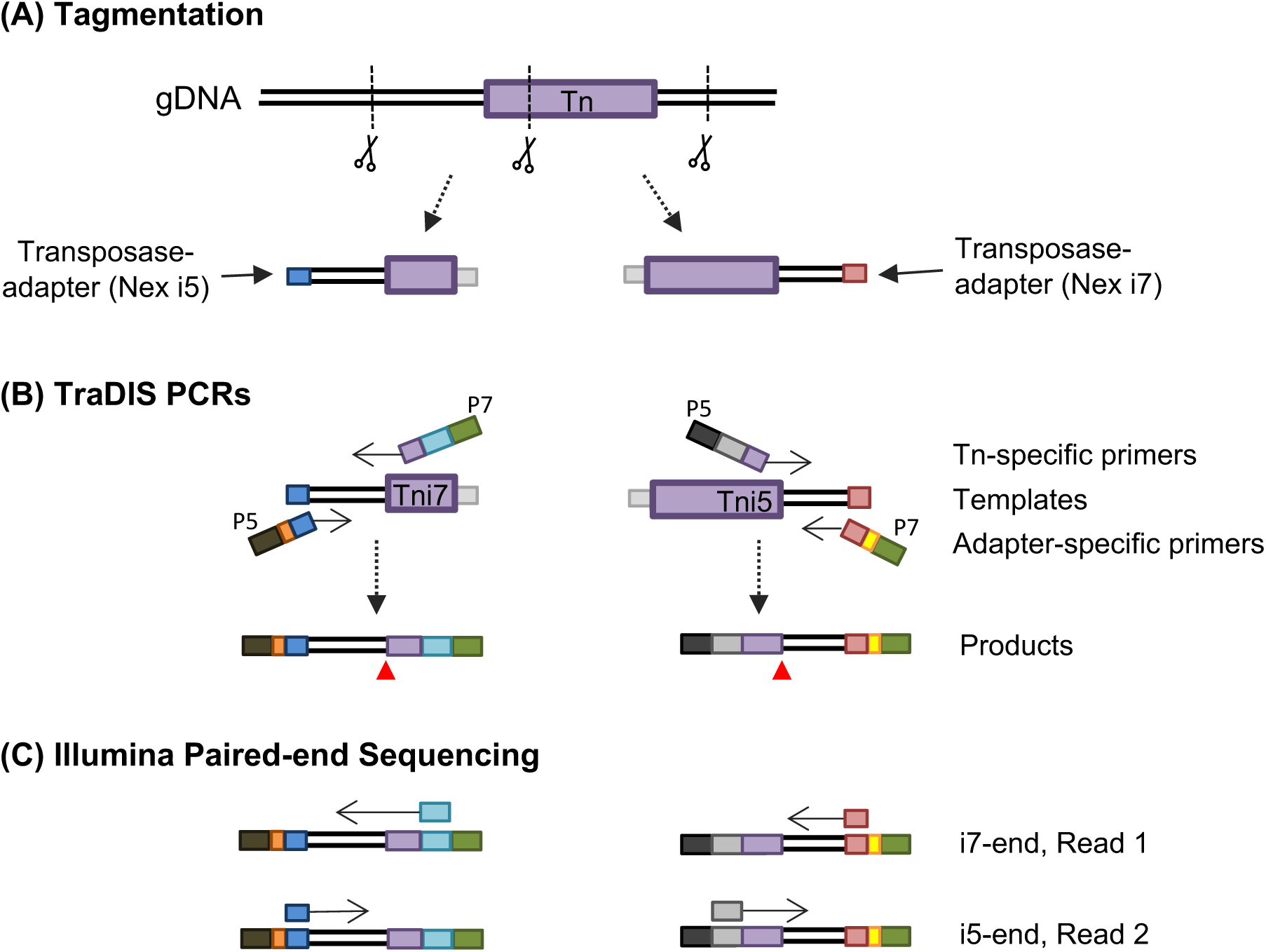
Outline of the modified transposon-directed insertion-site sequencing (TraDIS) protocol. **(A)** A genomic region containing a transposon (Tn) insertion site from the library is represented here. Genomic DNA (gDNA) is first randomly fragmented via tagmentation. In this reaction, the transposase cuts dsDNA and attaches standard Nextera-Illumina adapters (Nex i5, blue, or Nex i7, red) to the ends of the fragmented DNA. **(B)** Each end of the transposon is designated (“Tni5” and “Tni7”) by respective Tn-specific outward-directed primers that incorporate the appropriate 5’ Illumina flow-cell adapter (P5 or P7; dark grey and dark green, respectively) and read primer binding site (read 1 – cyan, and read 2 – light grey). For PCR, the opposing primer anneals to the Nex i5 or Nex i7 adapters (blue and red, respectively). These primers also contain unique barcodes to identify each sample (orange and yellow), and the alternate flow cell binding sequences (dark grey or dark green). The TraDIS PCR amplifies the transposon-gDNA junctions (indicated by red arrow-heads in the PCR products). **(C)** Paired-end sequencing is then carried out with standard Illumina workflows; multiplex sequencing of samples amplified from both ends of the transposon ensures high sequence diversity during both read 1 and read 2. For data analysis, high-quality sequence reads containing the expected transposon end sequence are selected, precisely identifying the collection of insertion sites.

**Supplementary Data Figure 2.**
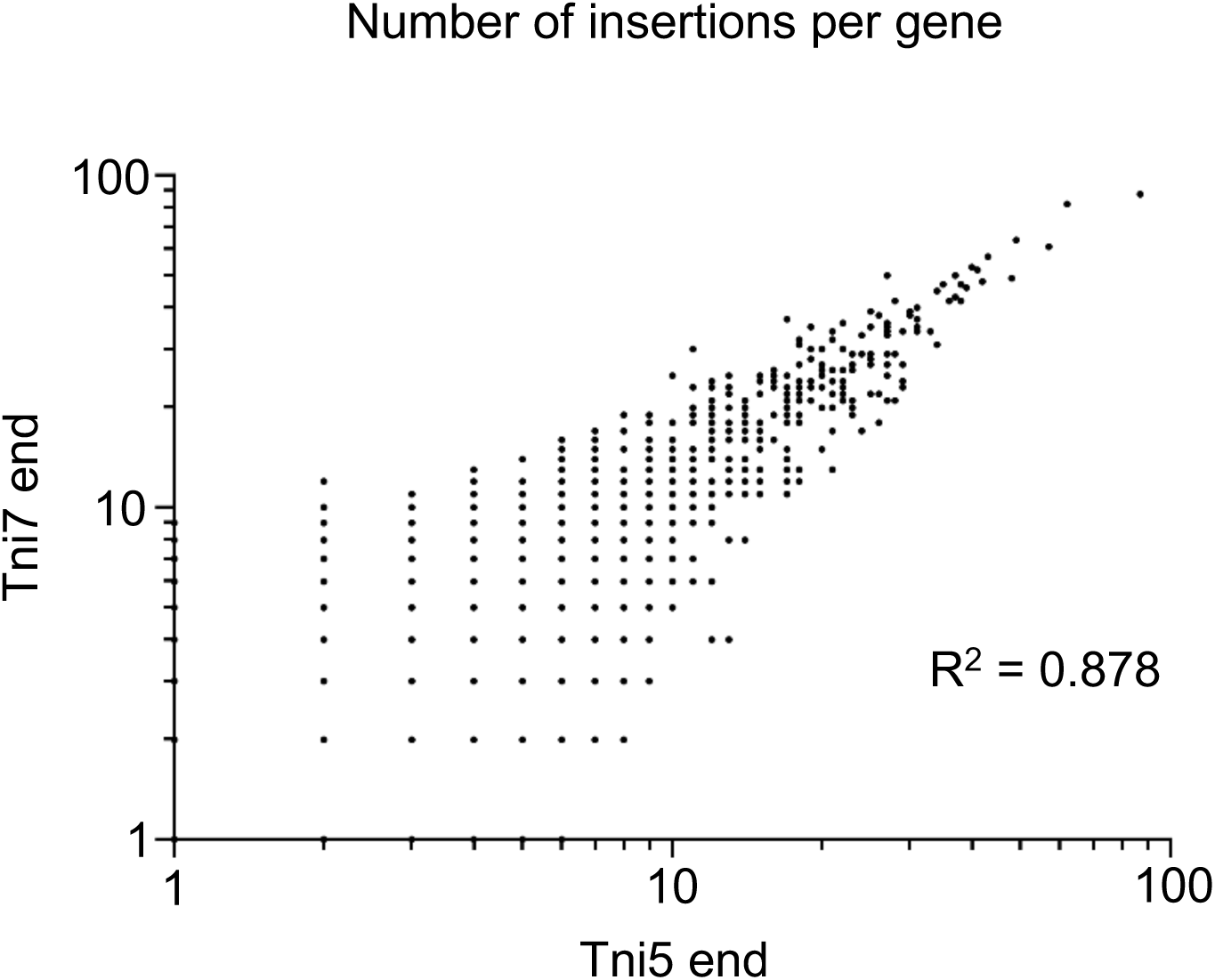
Correlation between the Tni5 and Tni7 sequencing ends of the *E. coli* UTI89 mini-Tn5 library. The number of insertion sites per gene in UTI89 was determined by sequencing the Tn-gDNA junctions from both the designated “Tni5” and “Tni7” ends of the transposon, mapping these reads the the UTI89 genome, and then identifying the number of specific insertion sites for each gene. For each gene, the number of insertions detected from the Tni5 and Tni7 ends was plotted; note that most points represent many genes.

**Supplementary Data Figure 3.**
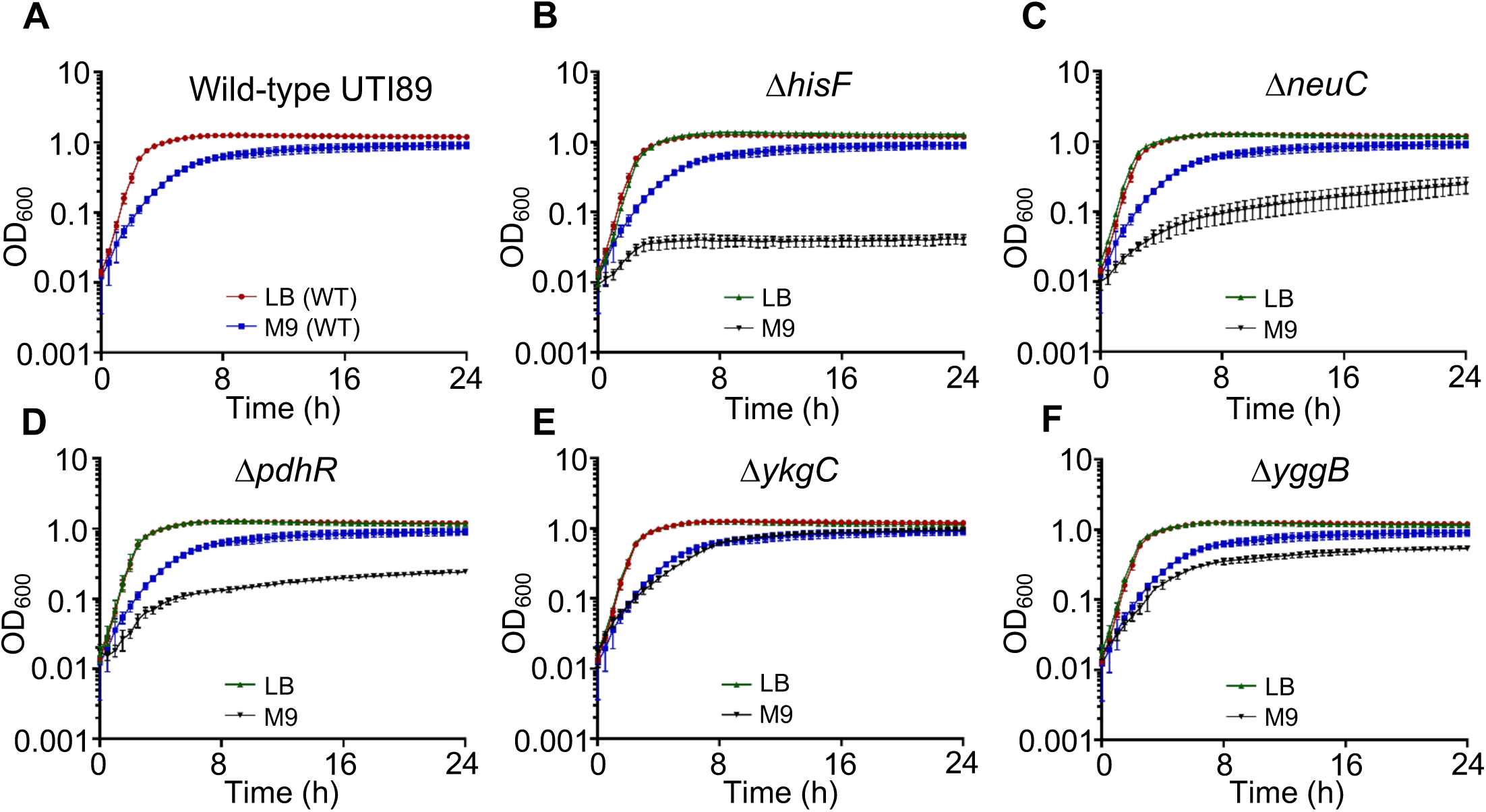
Growth curves of selected *E. coli* UTI89 gene deletion mutants. Wild-type UTI89 and the indicated deletion mutants grown in LB or M9-glycerol, as indicated. The wild-type UTI89 growth curves in LB (red) and M9-glycerol (blue) from (A) are also shown in these colors on the graphs for the deletions (B-F) for comparison.

**Supplementary Data Figure 4.**
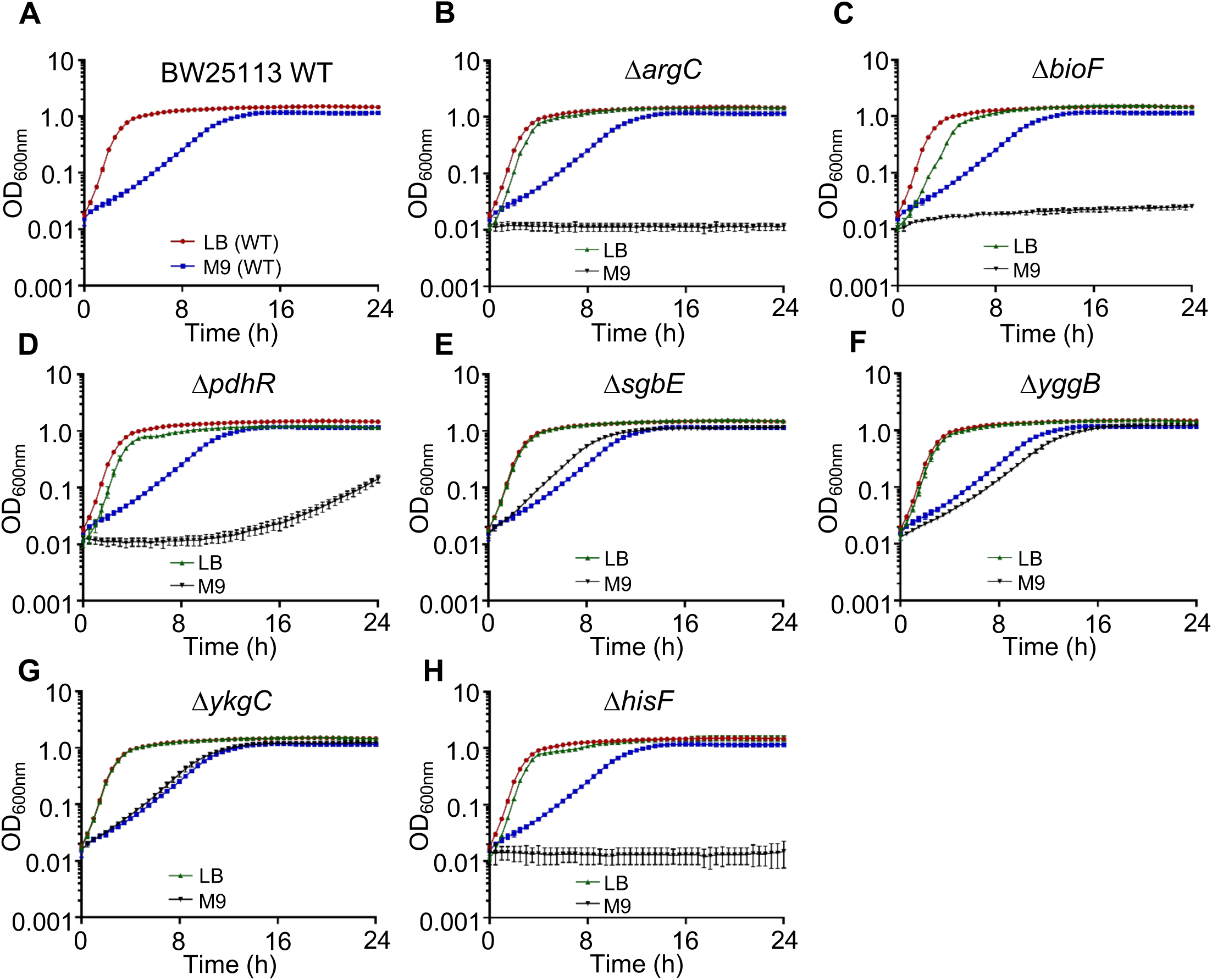
Growth curves of selected *E. coli* K-12 (BW25113) gene deletion mutants (from the KEIO collection). (A) BW25113 wild-type and the indicated deletion mutants grown in LB or M9-glycerol, as indicated. The wild-type BW25113 growth curves in LB (red) and M9-glycerol (blue) from (A) are also shown in these colors (B-H) for comparison.

**Supplementary Data Figure 5.**
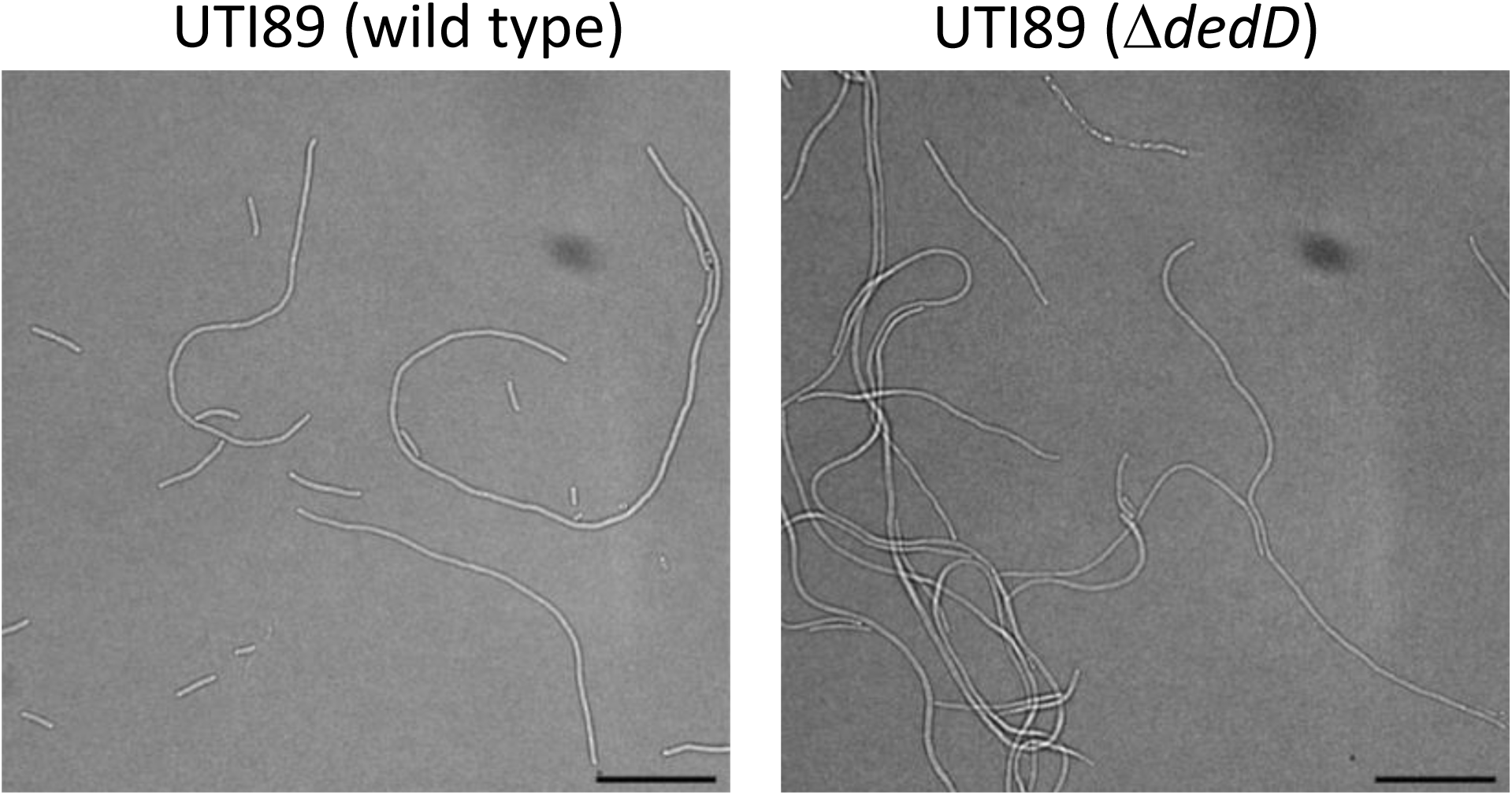
*dedD* deletion in UTI89 does not substantially affect infection-related filamentation (IRF) during dispersal from infection. UTI89 wild-type and 11*dedD* strains were sampled at dispersal stage of infection and then imaged by brightfield microscopy. Scale bars, 20 μm.

**Supplementary Data Figure 6.**
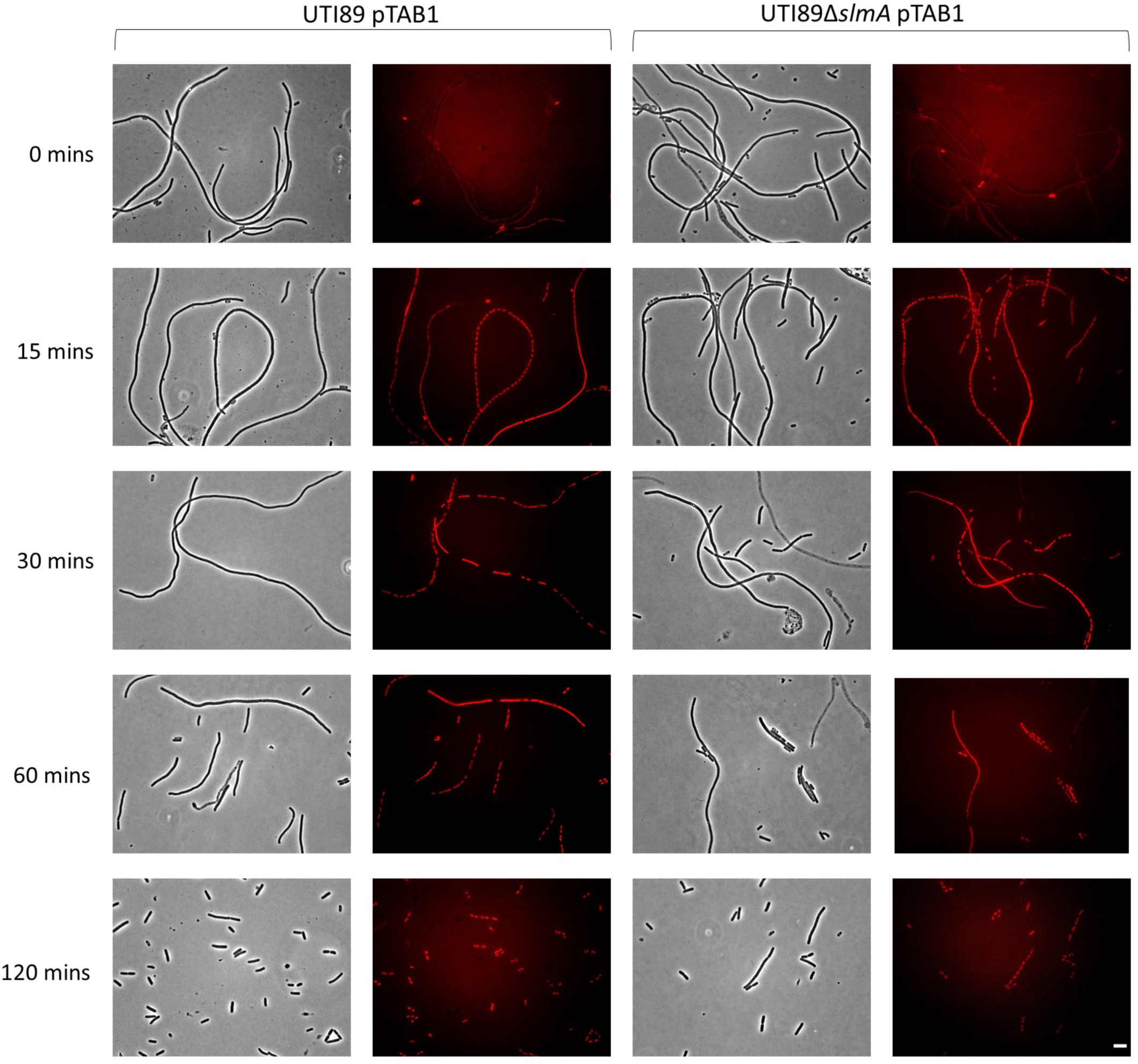
*slmA* deletion in UTI89 does not substantially affect filament reversal during recovery from infection. Wild-type and UTI89. 11*slmA* strains (containing an empty vector, pTAB1) were collected at dispersal stage of infections, and then grown in liquid LB culture and sampled at the indicated times for phase contrast (left) and fluorescence (right) microscopy – cells were fixed on sampling with 4% paraformaldehyde and then stained with 1 μg/mL DAPI DNA stain prior to microscopy. By 120 min, filaments of both strains had undergone substantial reversal into shorter rods. Scale bar, 5 μm (applicable to all images). Similar results were seen with time-lapse imaging of filament reversal on LB agar in situ (Supplementary Movie 1).

## REFERENCES

1. Lewis AJ, Richards AC, Mulvey MA. Invasion of Host Cells and Tissues by Uropathogenic Bacteria. Microbiol Spectr. 2016;4(6). doi: 10.1128/microbiolspec.UTI-0026-2016. PubMed PMID: 28087946; PubMed Central PMCID: PMC5244466.

2. Mulvey MA, Schilling JD, Hultgren SJ. Establishment of a persistent Escherichia coli reservoir during the acute phase of a bladder infection. Infection and immunity. 2001;69(7):4572–9. Epub 2001/06/13. doi: 10.1128/iai.69.7.4572-4579.2001. PubMed PMID: 11402001; PubMed Central PMCID: PMC98534.

3. Iosifidis G, Duggin IG. Distinct Morphological Fates of Uropathogenic Escherichia coli Intracellular Bacterial Communities: Dependency on Urine Composition and pH. Infect Immun. 2020;88(9). Epub 2020/06/17. doi: 10.1128/IAI.00884-19. PubMed PMID: 32540870; PubMed Central PMCID: PMCPMC7440767.

4. Justice SS, Hung C, Theriot JA, Fletcher DA, Anderson GG, Footer MJ, et al. Differentiation and developmental pathways of uropathogenic Escherichia coli in urinary tract pathogenesis. Proceedings of the National Academy of Sciences of the United States of America. 2004;101(5):1333–8. Epub 2004/01/24. doi: 10.1073/pnas.0308125100. PubMed PMID: 14739341; PubMed Central PMCID: PMC337053.

5. Mann R, Mediati DG, Duggin IG, Harry EJ, Bottomley AL. Metabolic Adaptations of Uropathogenic E. coli in the Urinary Tract. Frontiers in cellular and infection microbiology. 2017;7:241. doi: 10.3389/fcimb.2017.00241. PubMed PMID: 28642845; PubMed Central PMCID: PMC5463501.

6. Gerdes SY, Scholle MD, Campbell JW, Balazsi G, Ravasz E, Daugherty MD, et al. Experimental determination and system level analysis of essential genes in Escherichia coli MG1655. Journal of bacteriology. 2003;185(19):5673–84. Epub 2003/09/18. PubMed PMID: 13129938; PubMed Central PMCID: PMC193955.

7. Joyce AR, Reed JL, White A, Edwards R, Osterman A, Baba T, et al. Experimental and computational assessment of conditionally essential genes in Escherichia coli. J Bacteriol. 2006;188(23):8259–71. doi: 10.1128/JB.00740-06. PubMed PMID: 17012394; PubMed Central PMCID: PMC1698209.

8. Cain AK, Barquist L, Goodman AL, Paulsen IT, Parkhill J, van Opijnen T. A decade of advances in transposon-insertion sequencing. Nat Rev Genet. 2020;21(9):526–40. Epub 2020/06/14. doi: 10.1038/s41576-020-0244-x. PubMed PMID: 32533119; PubMed Central PMCID: PMCPMC7291929.

9. Goodall ECA, Robinson A, Johnston IG, Jabbari S, Turner KA, Cunningham AF, et al. The Essential Genome of Escherichia coli K-12. mBio. 2018;9(1). Epub 2018/02/22. doi: 10.1128/mBio.02096-17. PubMed PMID: 29463657; PubMed Central PMCID: PMCPMC5821084.

10. Langridge GC, Phan MD, Turner DJ, Perkins TT, Parts L, Haase J, et al. Simultaneous assay of every Salmonella Typhi gene using one million transposon mutants. Genome Res. 2009;19(12):2308–16. Epub 2009/10/15. doi: 10.1101/gr.097097.109. PubMed PMID: 19826075; PubMed Central PMCID: PMC2792183.

11. Phan MD, Peters KM, Sarkar S, Lukowski SW, Allsopp LP, Gomes Moriel D, et al. The serum resistome of a globally disseminated multidrug resistant uropathogenic Escherichia coli clone. PLoS Genet. 2013;9(10):e1003834. doi: 10.1371/journal.pgen.1003834. PubMed PMID: 24098145; PubMed Central PMCID: PMC3789825.

12. Goh KGK, Phan MD, Forde BM, Chong TM, Yin WF, Chan KG, et al. Genome-Wide Discovery of Genes Required for Capsule Production by Uropathogenic Escherichia coli. mBio. 2017;8(5). Epub 2017/10/27. doi: 10.1128/mBio.01558-17. PubMed PMID: 29066548; PubMed Central PMCID: PMCPMC5654933.

13. Sharp C, Boinett C, Cain A, Housden NG, Kumar S, Turner K, et al. O-Antigen-Dependent Colicin Insensitivity of Uropathogenic Escherichia coli. J Bacteriol. 2019;201(4). Epub 2018/12/05. doi: 10.1128/JB.00545-18. PubMed PMID: 30510143; PubMed Central PMCID: PMCPMC6351738.

14. Turner AK, Yasir M, Bastkowski S, Telatin A, Page AJ, Charles IG, et al. A genome-wide analysis of Escherichia coli responses to fosfomycin using TraDIS-Xpress reveals novel roles for phosphonate degradation and phosphate transport systems. J Antimicrob Chemother. 2020;75(11):3144–51. Epub 2020/08/07. doi: 10.1093/jac/dkaa296. PubMed PMID: 32756955; PubMed Central PMCID: PMCPMC7566553.

15. Yasir M, Turner AK, Bastkowski S, Baker D, Page AJ, Telatin A, et al. TraDIS-Xpress: a high-resolution whole-genome assay identifies novel mechanisms of triclosan action and resistance. Genome Res. 2020;30(2):239–49. Epub 2020/02/14. doi: 10.1101/gr.254391.119. PubMed PMID: 32051187; PubMed Central PMCID: PMCPMC7050523.

16. Meeske AJ, Rodrigues CD, Brady J, Lim HC, Bernhardt TG, Rudner DZ. High-Throughput Genetic Screens Identify a Large and Diverse Collection of New Sporulation Genes in Bacillus subtilis. PLoS Biol. 2016;14(1):e1002341. Epub 2016/01/07. doi: 10.1371/journal.pbio.1002341. PubMed PMID: 26735940; PubMed Central PMCID: PMCPMC4703394.

17. Liu X, Kimmey JM, Matarazzo L, de Bakker V, Van Maele L, Sirard JC, et al. Exploration of Bacterial Bottlenecks and Streptococcus pneumoniae Pathogenesis by CRISPRi-Seq. Cell Host Microbe. 2021;29(1):107–20 e6. Epub 2020/10/30. doi: 10.1016/j.chom.2020.10.001. PubMed PMID: 33120116; PubMed Central PMCID: PMCPMC7855995.

18. Karlinsey JE, Stepien TA, Mayho M, Singletary LA, Bingham-Ramos LK, Brehm MA, et al. Genome-wide Analysis of Salmonella enterica serovar Typhi in Humanized Mice Reveals Key Virulence Features. Cell Host Microbe. 2019;26(3):426–34 e6. Epub 2019/08/27. doi: 10.1016/j.chom.2019.08.001. PubMed PMID: 31447308; PubMed Central PMCID: PMCPMC6742556.

19. McCarthy AJ, Stabler RA, Taylor PW. Genome-Wide Identification by Transposon Insertion Sequencing of Escherichia coli K1 Genes Essential for In Vitro Growth, Gastrointestinal Colonizing Capacity, and Survival in Serum. J Bacteriol. 2018;200(7). Epub 2018/01/18. doi: 10.1128/JB.00698-17. PubMed PMID: 29339415; PubMed Central PMCID: PMCPMC5847654.

20. Subashchandrabose S, Smith S, DeOrnellas V, Crepin S, Kole M, Zahdeh C, et al. Acinetobacter baumannii Genes Required for Bacterial Survival during Bloodstream Infection. mSphere. 2016;1(1). Epub 2016/06/16. doi: 10.1128/mSphere.00013-15. PubMed PMID: 27303682; PubMed Central PMCID: PMCPMC4863628.

21. Subashchandrabose S, Smith SN, Spurbeck RR, Kole MM, Mobley HL. Genome-wide detection of fitness genes in uropathogenic Escherichia coli during systemic infection. PLoS Pathog. 2013;9(12):e1003788. Epub 2013/12/18. doi: 10.1371/journal.ppat.1003788. PubMed PMID: 24339777; PubMed Central PMCID: PMCPMC3855560.

22. van Opijnen T, Camilli A. A fine scale phenotype-genotype virulence map of a bacterial pathogen. Genome Res. 2012;22(12):2541–51. Epub 2012/07/25. doi: 10.1101/gr.137430.112. PubMed PMID: 22826510; PubMed Central PMCID: PMCPMC3514683.

23. Garcia V, Gronnemose RB, Torres-Puig S, Kudirkiene E, Piantelli M, Ahmed S, et al. Genome-wide analysis of fitness-factors in uropathogenic Escherichia coli during growth in laboratory media and during urinary tract infections. Microb Genom. 2021;7(12). doi: 10.1099/mgen.0.000719. PubMed PMID: 34928200; PubMed Central PMCID: PMCPMC8767336.

24. Garcia V, Staerk K, Alobaidallah MSA, Gronnemose RB, Guerra PR, Andersen TE, et al. Genome-wide analysis of fitness factors in uropathogenic Escherichia coli in a pig urinary tract infection model. Microbiol Res. 2022;265:127202. Epub 20220915. doi: 10.1016/j.micres.2022.127202. PubMed PMID: 36167007.

25. Andersen TE, Khandige S, Madelung M, Brewer J, Kolmos HJ, Moller-Jensen J. Escherichia coli uropathogenesis in vitro: invasion, cellular escape, and secondary infection analyzed in a human bladder cell infection model. Infection and immunity. 2012;80(5):1858–67. Epub 2012/02/23. doi: 10.1128/iai.06075-11. PubMed PMID: 22354025; PubMed Central PMCID: PMC3347433.

26. Khandige S, Asferg CA, Rasmussen KJ, Larsen MJ, Overgaard M, Andersen TE, et al. DamX Controls Reversible Cell Morphology Switching in Uropathogenic Escherichia coli. mBio. 2016;7(4). doi: 10.1128/mBio.00642-16. PubMed PMID: 27486187; PubMed Central PMCID: PMC4981707.

27. Baba T, Ara T, Hasegawa M, Takai Y, Okumura Y, Baba M, et al. Construction of Escherichia coli K-12 in-frame, single-gene knockout mutants: the Keio collection. Mol Syst Biol. 2006;2:2006 0008. Epub 2006/06/02. doi: msb4100050 [pii] 10.1038/msb4100050. PubMed PMID: 16738554; PubMed Central PMCID: PMC1681482.

28. Dembek M, Barquist L, Boinett CJ, Cain AK, Mayho M, Lawley TD, et al. High-throughput analysis of gene essentiality and sporulation in Clostridium difficile. mBio. 2015;6(2):e02383. doi: 10.1128/mBio.02383-14. PubMed PMID: 25714712; PubMed Central PMCID: PMC4358009.

29. Rosconi F, Rudmann E, Li J, Surujon D, Anthony J, Frank M, et al. A bacterial pan-genome makes gene essentiality strain-dependent and evolvable. Nat Microbiol. 2022;7(10):1580–92. Epub 20220912. doi: 10.1038/s41564-022-01208-7. PubMed PMID: 36097170; PubMed Central PMCID: PMCPMC9519441.

30. Rosen DA, Hooton TM, Stamm WE, Humphrey PA, Hultgren SJ. Detection of intracellular bacterial communities in human urinary tract infection. PLoS Med. 2007;4(12):e329. Epub 2007/12/21. doi: 10.1371/journal.pmed.0040329. PubMed PMID: 18092884; PubMed Central PMCID: PMC2140087.

31. Abell-King C, Costas A, Duggin IG, Soderstrom B. Bacterial filamentation during urinary tract infections. PLoS Pathog. 2022;18(12):e1010950. Epub 20221201. doi: 10.1371/journal.ppat.1010950. PubMed PMID: 36454736; PubMed Central PMCID: PMCPMC9714745.

32. Soderstrom B, Pittorino MJ, Daley DO, Duggin IG. Assembly dynamics of FtsZ and DamX during infection-related filamentation and division in uropathogenic E. coli. Nat Commun. 2022;13(1):3648. Epub 20220625. doi: 10.1038/s41467-022-31378-1. PubMed PMID: 35752634; PubMed Central PMCID: PMCPMC9233674.

33. Raterman EL, Shapiro DD, Stevens DJ, Schwartz KJ, Welch RA. Genetic analysis of the role of yfiR in the ability of Escherichia coli CFT073 to control cellular cyclic dimeric GMP levels and to persist in the urinary tract. Infect Immun. 2013;81(9):3089–98. Epub 20130617. doi: 10.1128/IAI.01396-12. PubMed PMID: 23774594; PubMed Central PMCID: PMCPMC3754225.

34. Kim HK, Harshey RM. A Diguanylate Cyclase Acts as a Cell Division Inhibitor in a Two-Step Response to Reductive and Envelope Stresses. mBio. 2016;7(4). Epub 2016/08/11. doi: 10.1128/mBio.00822-16. PubMed PMID: 27507823; PubMed Central PMCID: PMCPMC4992967.

35. Bernhardt TG, de Boer PA. SlmA, a nucleoid-associated, FtsZ binding protein required for blocking septal ring assembly over Chromosomes in E. coli. Molecular cell. 2005;18(5):555–64. doi: 10.1016/j.molcel.2005.04.012. PubMed PMID: 15916962; PubMed Central PMCID: PMC4428309.

36. Pagnout C, Sohm B, Razafitianamaharavo A, Caillet C, Offroy M, Leduc M, et al. Pleiotropic effects of rfa-gene mutations on Escherichia coli envelope properties. Sci Rep. 2019;9(1):9696. Epub 20190704. doi: 10.1038/s41598-019-46100-3. PubMed PMID: 31273247; PubMed Central PMCID: PMCPMC6609704.

37. Alteri CJ, Mobley HL. Metabolism and Fitness of Urinary Tract Pathogens. Microbiol Spectr. 2015;3(3). doi: 10.1128/microbiolspec.MBP-0016-2015. PubMed PMID: 26185076; PubMed Central PMCID: PMC4510461.

38. Alteri CJ, Smith SN, Mobley HL. Fitness of Escherichia coli during urinary tract infection requires gluconeogenesis and the TCA cycle. PLoS Pathog. 2009;5(5):e1000448. doi: 10.1371/journal.ppat.1000448. PubMed PMID: 19478872; PubMed Central PMCID: PMC2680622.

39. Conover MS, Hadjifrangiskou M, Palermo JJ, Hibbing ME, Dodson KW, Hultgren SJ. Metabolic Requirements of Escherichia coli in Intracellular Bacterial Communities during Urinary Tract Infection Pathogenesis. mBio. 2016;7(2):e00104–16. doi: 10.1128/mBio.00104-16. PubMed PMID: 27073089; PubMed Central PMCID: PMC4959519.

40. Vann WF, Daines DA, Murkin AS, Tanner ME, Chaffin DO, Rubens CE, et al. The NeuC protein of Escherichia coli K1 is a UDP N-acetylglucosamine 2-epimerase. J Bacteriol. 2004;186(3):706–12. Epub 2004/01/20. doi: 10.1128/jb.186.3.706-712.2004. PubMed PMID: 14729696; PubMed Central PMCID: PMCPMC321479.

41. Anderson GG, Goller CC, Justice S, Hultgren SJ, Seed PC. Polysaccharide capsule and sialic acid-mediated regulation promote biofilm-like intracellular bacterial communities during cystitis. Infect Immun. 2010;78(3):963–75. Epub 2010/01/21. doi: 10.1128/IAI.00925-09. PubMed PMID: 20086090; PubMed Central PMCID: PMCPMC2825929.

42. Burns SM, Hull SI. Comparison of loss of serum resistance by defined lipopolysaccharide mutants and an acapsular mutant of uropathogenic Escherichia coli O75:K5. Infect Immun. 1998;66(9):4244–53. doi: 10.1128/IAI.66.9.4244-4253.1998. PubMed PMID: 9712774; PubMed Central PMCID: PMCPMC108512.

43. Burns SM, Hull SI. Loss of resistance to ingestion and phagocytic killing by O(-) and K(-) mutants of a uropathogenic Escherichia coli O75:K5 strain. Infect Immun. 1999;67(8):3757–62. doi: 10.1128/IAI.67.8.3757-3762.1999. PubMed PMID: 10417134; PubMed Central PMCID: PMCPMC96650.

44. Sarkar S, Ulett GC, Totsika M, Phan MD, Schembri MA. Role of capsule and O antigen in the virulence of uropathogenic Escherichia coli. PLoS One. 2014;9(4):e94786. Epub 2014/04/12. doi: 10.1371/journal.pone.0094786. PubMed PMID: 24722484; PubMed Central PMCID: PMCPMC3983267.

45. Sellner B, Prakapaite R, van Berkum M, Heinemann M, Harms A, Jenal U. A New Sugar for an Old Phage: a c-di-GMP-Dependent Polysaccharide Pathway Sensitizes Escherichia coli for Bacteriophage Infection. mBio. 2021;12(6):e0324621. Epub 20211214. doi: 10.1128/mbio.03246-21. PubMed PMID: 34903045; PubMed Central PMCID: PMCPMC8669472.

46. Klein K, Palarasah Y, Kolmos HJ, Moller-Jensen J, Andersen TE. Quantification of filamentation by uropathogenic Escherichia coli during experimental bladder cell infection by using semi-automated image analysis. J Microbiol Methods. 2015;109:110–6. Epub 2014/12/30. doi: 10.1016/j.mimet.2014.12.017. PubMed PMID: 25546841.

47. Bottomley AL, Peterson E, Iosifidis G, Yong AMH, Hartley-Tassell LE, Ansari S, et al. The novel E. coli cell division protein, YtfB, plays a role in eukaryotic cell adhesion. Sci Rep. 2020;10(1):6745. Epub 2020/04/23. doi: 10.1038/s41598-020-63729-7. PubMed PMID: 32317661; PubMed Central PMCID: PMCPMC7174318.

48. Jorgenson MA, Young KD. YtfB, an OapA domain-containing protein, is a new cell division protein in Escherichia coli. J Bacteriol. 2018. Epub 2018/04/25. doi: 10.1128/JB.00046-18. PubMed PMID: 29686141; PubMed Central PMCID: PMCPMC5996693.

49. Liu B, Hale CA, Persons L, Phillips-Mason PJ, de Boer PAJ. Roles of the DedD Protein in Escherichia coli Cell Constriction. J Bacteriol. 2019;201(8). Epub 20190326. doi: 10.1128/JB.00698-18. PubMed PMID: 30692172; PubMed Central PMCID: PMCPMC6436348.

50. Barquist L, Mayho M, Cummins C, Cain AK, Boinett CJ, Page AJ, et al. The TraDIS toolkit: sequencing and analysis for dense transposon mutant libraries. Bioinformatics. 2016;32(7):1109–11. doi: 10.1093/bioinformatics/btw022. PubMed PMID: 26794317; PubMed Central PMCID: PMC4896371.

51. Ramirez F, Ryan DP, Gruning B, Bhardwaj V, Kilpert F, Richter AS, et al. deepTools2: a next generation web server for deep-sequencing data analysis. Nucleic Acids Res. 2016;44(W1):W160–5. Epub 2016/04/16. doi: 10.1093/nar/gkw257. PubMed PMID: 27079975; PubMed Central PMCID: PMCPMC4987876.

52. Abd-El-Haliem A. An unbiased method for the quantitation of disease phenotypes using a custom-built macro plugin for the program ImageJ. 2012;835:635–44. doi: 10.1007/978-1-61779-501-5_41. PubMed PMID: 22183684.

53. Wright K, de Silva K, Plain KM, Purdie AC, Blair TA, Duggin IG, et al. Mycobacterial infection-induced miR-206 inhibits protective neutrophil recruitment via the CXCL12/CXCR4 signalling axis. PLoS Pathog. 2021;17(4):e1009186. Epub 20210407. doi: 10.1371/journal.ppat.1009186. PubMed PMID: 33826679; PubMed Central PMCID: PMCPMC8055004.

